# Maturation of Dorsal Association Tracts during Preadolescence Links to Concurrent and Future Cognitive Performance and Transdiagnostic Psychopathology

**DOI:** 10.1101/2025.07.17.664983

**Authors:** Danni Wang, Christopher J Hammond, Betty Jo Salmeron, Xiang Xiao, Laura Murray, Hong Gu, Tianye Zhai, Annika Quam, Justine Hill, Hieu Nguyen, Hanbing Lu, Amy Janes, Thomas J Ross, Yihong Yang

## Abstract

Many psychiatric disorders begin during adolescence, coinciding with the rapid development of brain white matter (WM). However, it remains unclear whether deviations from normal WM maturation during this age period contribute to the development of psychopathology. In this study, we developed and validated normative models of brain age based on specific WM tracts using three large-scale developmental datasets (a total of ∼10,000 subjects). We found that tract-specific deviations in WM development of association and limbic/subcortical systems were linked to concurrent cognition and psychopathology. The spatial pattern of the association system aligned closely with distributions of high-order brain networks, and with mitochondrial content and respiratory capacity. The maturation of the association system contributed significantly to better cognitive performance assessed two or three years later. Importantly, delayed WM development especially in dorsal association tracts predicted psychiatric disorders across diagnoses and disorder onset over a 2-year follow-up. By identifying tract-specific WM development during preadolescence as a predictor of cognitive capacity and psychiatric disorder risks, this study provides a valuable framework for tracking individualized brain maturation and understanding the neurobiological underpinnings of cognitive performance and transdiagnostic psychopathology.

## Introduction

Psychiatric disorders are among the most common causes of morbidity and mortality ^1, 2^, yet much is still unknown regarding their underlying developmental pathophysiologies^3^. One emerging hypothesis is that a large amount of variance in the risk for psychopathology comes from common “transdiagnostic” instead of disorder-specific factors. This is supported by findings showing that different psychiatric disorders share risk genes and show overlapping alterations in brain structure, connectivity and function^4, 5, 6^. Further, approximately 75% of all psychiatric disorders start before the age of 21, with 35% starting before age of 14^7^, suggesting that deviations in neurodevelopment may contribute to psychopathology risk in a transdiagnostic manner.

Personalized characterization of white matter (WM) tract development may provide significant insight into links between disrupted neurodevelopment and psychopathology as there is significant temporal overlap between WM changes during preadolescence and the onset of many psychiatric disorders^8, 9, 10^. It is plausible that WM alterations may contribute to mental health issues given that WM tract integrity in association and limbic tracts are linked with cognitive and socioemotional processes in pediatric populations^11^. Developmental variance in the structural integrity of these critical pathways may have widespread influences on cognitive and emotional functioning, which may in turn influence risk for psychopathology^12, 13^. Further, the energetic demands of WM is higher during the preadolescent and early adolescent window, driven by maturation-related increases in physiological processes such as myelin synthesis and axon– oligodendrocyte^9^. Mitochondria, which are central to energy metabolism, have been shown in histological studies to exhibit substantial alterations in individuals with psychiatric disorders^14, 15^, implicating mitochondrial dysfunction as a potential contributor to impaired WM integrity. Given that WM enables efficient inter-regional communication and supports the transmission of neurotransmitter signals, its developmental disruption could have cascading effects on brain function. However, links between WM development, mitochondrial functioning, and psychopathology are poorly understand.

One way to assess normative growth and define pathological deviations in development is through the use of normative modeling which has been successfully applied in clinical contexts including growth charts in pediatrics^16^ and standardized achievement and intelligence (IQ) tests in psychology^17^. Recently, this framework has been extended to measures of brain structure and function to quantify person-specific deviations in developmental trajectories relative to typically developing controls^18, 19, 20, 21^. In the context of assessing individual brain development/aging level, machine learning prediction modeling using magnetic resonance imaging (MRI) data has allowed for the estimation of the biological age of the brain of healthy individuals^22^. Deviations between this neuroimaging-based ‘brain age’ and chronological age, termed the brain age gap (BAG), may reflect disruptions in normal developmental/aging trajectories and provide mechanistic information into cognitive development and the pathophysiology of psychiatric disorders^22, 23, 24^. To date, this method has focused on morphology studies using measures of gray matter (GM) volume, thickness, and area and such studies suggest that psychopathology is associated with accelerated brain aging in adults and youth^22, 23, 24^. While these prior studies highlight the relevance of the GM-derived BAG in psychopathology, the role of WM maturation is still unclear. Our study seeks to fill this knowledge gap.

In the present study, we sought to characterize WM BAG patterns derived from tract-specific microstructure in preadolescents with varying levels of psychopathology and examine associations between tract-based BAGs and a broad set of behavioral assessments spanning cognitive and psychopathology domains in youth. First, we built and cross-validated WM tract-based brain age models using diffusion MRI (dMRI) data from the Lifespan Human Connectome Project Development (HCP-D) dataset (*N* = 611). We then performed out-of-sample validation, using models derived from HCP-D data to predict brain ages in two independent datasets including Adolescent Brain Cognitive Development (ABCD) study (*N*_𝐵𝑎𝑠𝑒𝑙𝑖𝑛𝑒_ = 8,688) and Healthy Brain Network (HBN) pediatric mental health study (*N* = 978). Next, we used baseline data from the ABCD study and applied a multivariate sparse canonical correlation analysis (sCCA) to identify latent brain-behavior associations between WM-derived BAGs and cognition and psychopathology. Then, we assessed whether the tract-wise patterns of developmental levels linking cognition and psychopathology domains can be explained by tract-wise mitochondrial profiles from postmortem data^25^. Lastly, to ascertain the clinical utility of WM BAGs in follow-up cognitive performance and risk for psychiatric illness, we examined tract-based BAG associations with i) the cognitive performance measured 2 or 3 years after baseline, and ii) the cumulative number of psychiatric diagnoses assessed concurrently and 2 years after baseline, as well as longitudinal transitions between healthy and psychiatrically diagnosed states over the 2- year study period.

## Results

### Whole-brain and individual WM tract profiles predicted chronological age

First, we extracted 54 WM tracts from dMRI in three datasets (HCP-D, HBN, and ABCD) and categorized the tracts into 7 systems (Figure 1A). The GM regions connected by each tract are indicated in Table S1. Tract-based features were constructed using diffusion fractional anisotropy (FA) profiles along tracts (Figure 1B). We then built brain age models using these tract-based features of individual tracts (25 combined bilateral association/projection tracts and 4 callosal tracts) or whole brain (concatenated all tracts) from typically developing adolescents in HCP-D dataset (*N* = 611; 330 females, age range = 5.58-21.92 years) (Figure 1C). The models were trained by a Gaussian Process Regression (GPR) algorithm^26^, which utilizes a Bayesian approach to predict continuous variables and has shown strong predictive performance for age from MRI-derived metrics in previous studies^27, 28^. Performance of the brain age model from whole-brain tracts is shown in Figure S1. The 5-fold cross-validation performance of the GPR model showed a R^2^ of 0.702 and a mean absolute error (MAE) of 1.805 years. To test the prediction performance of the brain age model, we selected 978 participants from the independent HBN dataset (366 females, age range = 5.58-21.90 years) within the same age range as the HCP-D study. As shown in Table S2, whole-brain brain-age model significantly estimated age with R^2^ of 0.387 and MAE of 2.798 years. For the prediction of brain ages in another independent dataset, the ABCD study (*N*_𝐵𝑎𝑠𝑒𝑙𝑖𝑛𝑒_ = 8,688, 4,205 females, age range = 8.91-11.08 years; *N*_2–𝑦–𝑓𝑜𝑙𝑙𝑜w–𝑢𝑝_ = 5,883, 2,741 females; age range = 10.58-13.83 years), the age range was much narrower and the prediction performance was with a R^2^ of 0.185 and an MAE of 1.68 years (Figure S1).

**Figure 1.**
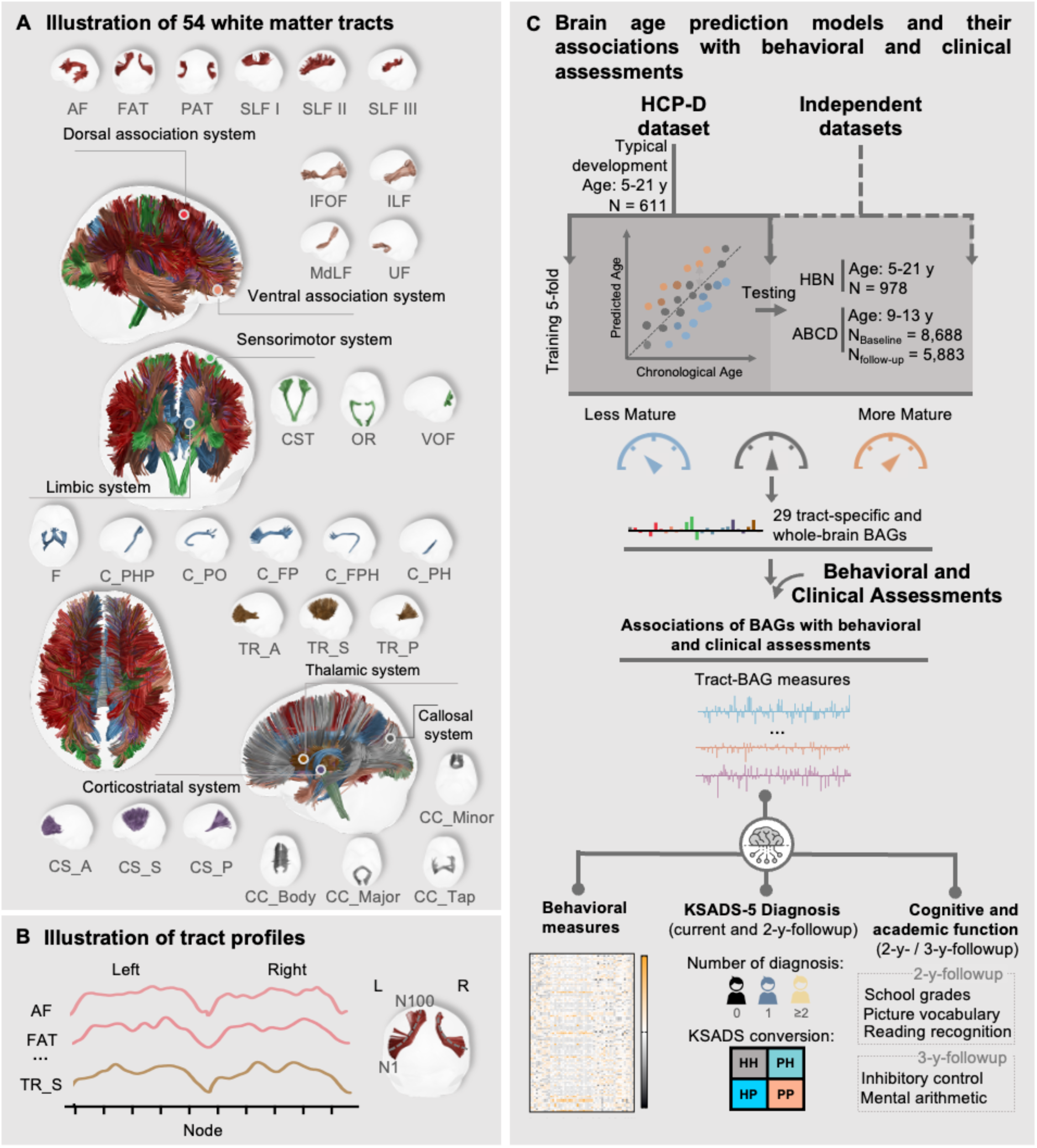
Overview of study design. (A) White matter tracts used for brain age modeling: 54 white matter tracts grouped into 7 distinct systems, for which normative models of brain age were constructed using tract-specific features. (B) Tract profiles of fractional anisotropy (FA). 100 segments (nodes) were evenly sampled along each unilateral tract, generating 200 FA values per bilateral tract. (C) Predictive models of brain age were generated using FA tract-profiles of the specific tracts from the participants in Human Connectome Project in Development (HCP-D) dataset with 5-fold cross-validation. One whole-brain and 29 specific-tract normative developmental models were established separately. Using models trained on HCP-D dataset, brain age gaps (BAGs) based on whole-brain and individual tracts were computed for each participant in the independent Healthy Brain Network pediatric mental health (HBN) and Adolescent Brain Cognitive Development (ABCD) datasets. Associations of the BAGs with behavioral and clinical assessments were investigated. The full names of the tracts are listed in the Table S1. Abbreviations: HH, healthy-persistent; PH, disorder-remitted; HP, disorder-new-onset; PP, disorder-persistent; KSADS-5, Kiddie Schedule for Affective Disorders and Schizophrenia for DSM-5.

For brain age models built from FA profiles of individual tracts, all 29 tract-based brain age models significantly predicted the age in the 5-fold cross-validation dataset (HCP-D) after false discovery rate (FDR) correction for multiple comparisons. The cross-validation performance in HCP-D dataset was with R^2^s ranging from 0.240 to 0.576 (MAEs ranging from 2.09 to 2.87). See Table S2 for the cross-validation performance of HCP-D dataset. For the prediction of brain age in testing datasets (HBN and ABCD), all tracts also exhibited significant prediction for age (Figure S1 and Tables S3-4).

For each participant, the WM tract predicted age is referred to as brain age, and the difference between the brain age and chronological age is defined as tract-based BAG. This tract-based BAG for an individual reflects whether the development of the tract appears less or more mature than expected relative to his/her chronological age. A negative BAG indicates that an individual’s predicted brain age is less than their chronological age (i.e., WM tract is less mature compared to age-matched normative model), while a positive BAG indicates that an individual’s predicted brain age is higher than their chronological age (i.e., WM tract is more mature compared to age-matched normative model). To correct the potential brain age bias caused by regression dilution, we further trained brain age models with bias correction, a linear transformation of chronological age^29^, on the training sets and applied them to the testing data sets. The predictive performance of the corrected tract-based brain ages is reported in Table S2-4 and Figure S1. We included both tract-based BAGs with or without age-bias correction in our statistical analyses. While there is ongoing debate regarding the optimal method for bias correction in brain age modeling^30, 31^, our key findings remained robust regardless of whether age-bias correction was applied, as the BAGs were only adjusted by a linear transformation of chronological age, and age was also included as a covariate in all subsequent analyses.

### Tract-based BAGs associated with cognition and psychopathology

To comprehensively characterize tract-based BAGs and their associations with general cognition and psychopathology, we examined associations of tract-based BAGs with a wide range of cognitive functions and psychopathological behaviors using multivariate sCCA, with age and sex as covariates. In the sCCA, we included 30 tract-based BAGs (29 tract-specific and a whole-brain tract BAGs) and 51 “behavioral assessments” (20 neurocognitive and 31 psychopathology-related measures, see Methods) from the baseline ABCD dataset, which were utilized in our previous functional study^32^. The analysis showed that the first two canonical modes were statistically significant compared to null distributions generated by permutation tests (sCCA mode 1: 𝑅 = 0.19, *p_FDR_* < 0.001 ; sCCA mode 2: 𝑅 = 0.13, *p_FDR_* < 0.001 ). The scree plot of the covariance explained by all canonical modes is shown in Figure S2.

The loading patterns of the tract-based BAGs on the two significant sCCA modes were highly heterogeneous (Figure 2A). BAGs obtained from most tracts in the dorsal and ventral association systems exhibited significant positive loadings for Mode 1. In contrast, BAGs from tracts mainly from the sensorimotor, limbic, cortico-striatum, cortico-thalamus and callosal systems had significant positive loadings for Mode 2. Therefore, in the following discussion, the 1^st^ sCCA mode is referred to as Association Development mode and the 2^nd^ mode is referred to as Subcortical/limbic Development mode.

**Figure 2.**
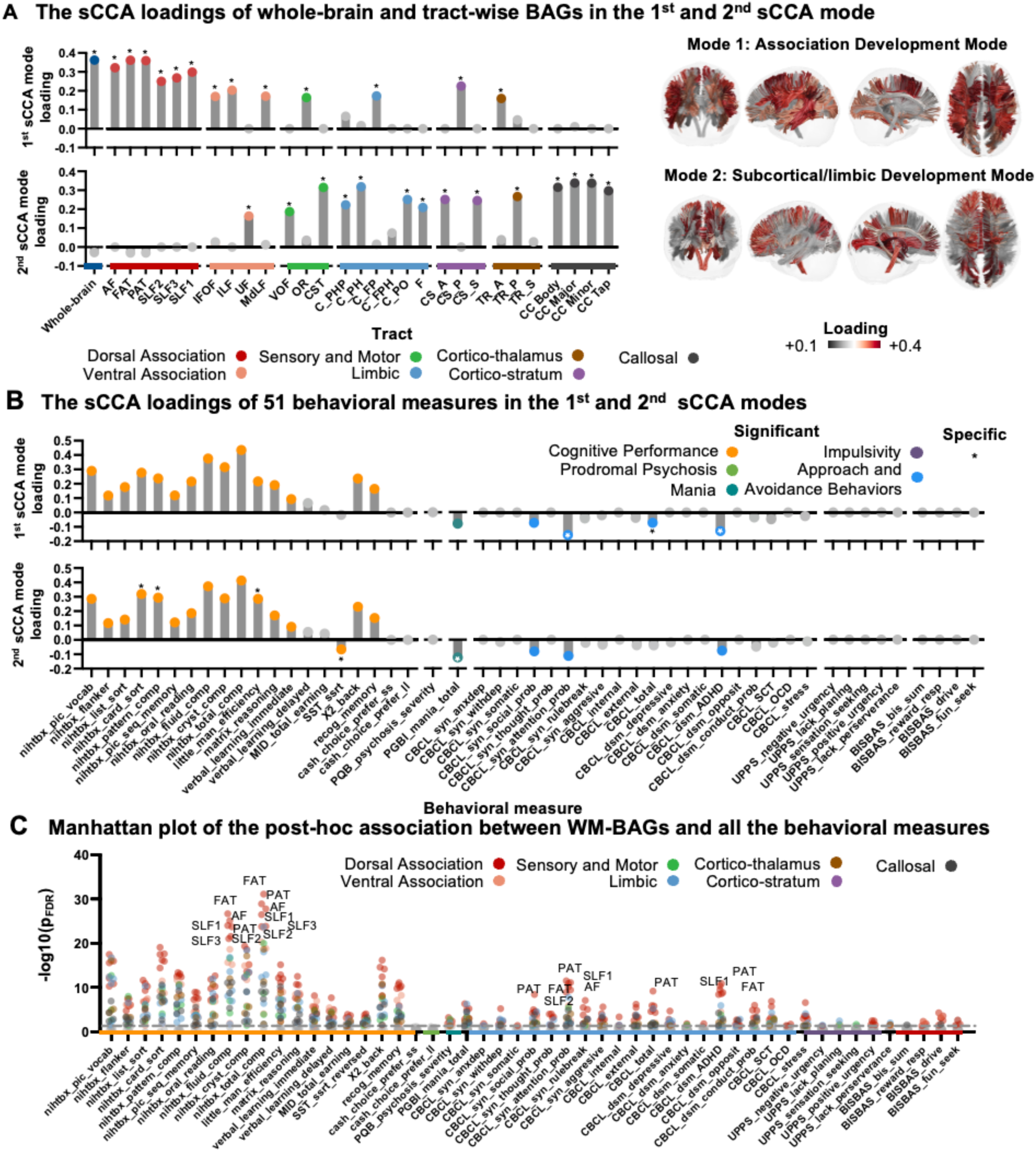
Multivariate associations between tract-based brain age gaps (BAGs) and behavioral assessments using sparse canonical correlation analysis (sCCA). The first two canonical variates were statistically significant, as determined by permutation testing with false discovery rate (FDR) correction (p < 0.05). (A) Loadings of the brain features on the two sCCA modes. (B) Loadings of behavioral assessments on the two sCCA modes. Statistically significant loadings are colored according to the behavioral/tract systems to which they belonged, with those specific to each mode highlighted with asterisks. (C) The Manhattan plot of the post-hoc associations between tract-based BAGs and all the behavioral measures. The dots above the gray dotted line indicate that the tract-based BAGs were significantly correlated with behavioral assessments (*p_FDR_* < 0.05). Tract-based BAGs with significant associations are labeled with colors. The top 10 tract-based BAGs that were significantly associated with cognition or psychopathology domains are highlighted with the tract names, respectively. The full names of the tracts are listed in the Table S1.

Cognitive measures had generally heavy and positive loadings on the two sCCA modes (Figure 2B). To evaluate the specific behavioral measures associated with each sCCA mode, we compared the distributions of loadings generated from 1,000 bootstrap tests (see Methods). The measures with an effect size more than 0.5 were considered specific to the corresponding mode. As shown in Figure 2A and Table S5, the Association Development mode significantly loaded on cognitive measures including total cognition composite and fluid cognition composite scores. Several psychopathological assessments, including Child Behavioral Checklist (CBCL) Attention Problems, Attention Deficit / Hyperactivity Disorder, and Total Problems exhibited significant and specific negative loadings on the Association Development mode. For the Subcortical/limbic Development mode, significant and specific cognitive scores included Little Man Task efficiency, Pattern Comparison processing speed and Stop Signal Reaction (SST) time, while Parent General Behavior Inventory (PGBI) Total Score of Mania showed significant negative loading.

Furthermore, we performed a post-hoc statistical analysis to explore the univariate relationships between the 30 tract-based BAGs and 51 behavioral measures. The heatmap of the t values is in Figure S3-4, showing significant associations of nearly all the cognitive measures with tract-based BAGs. The Manhattan plot in Figure 2C visualizes the associations between all tract-based BAGs and behavioral measures. The top 10 tract-based BAGs showing the strongest correlations with either cognitive or psychopathology domains are highlighted. Across both domains, dorsal association tracts consistently ranked higher than tracts from other systems. Notably, the behavioral measures showing the strongest associations with dorsal association tracts included total intelligence, fluid intelligence, CBCL Total Problems, Attention Problems, and ADHD scores.

### Functional decoding based on spatial association between brain regions connected by WM tracts and task-related brain activation

WM tracts connect distinct brain functional units that correspond to relevant task-related brain activations. To further characterize functions of GM regions connected by the WM tracts with loadings in the two identified sCCA modes, we utilized brain activation maps in the Neurosynth meta-analysis based on 100 topics^33^. For the functional activation map of each topic, we calculated the brain activation level within the GM regions connected by each association/projection WM tract. The topics (Table S6) related to high-order cognitive ability, emotion and psychiatric disorders were selected to perform spatial correlation with the loading patterns of the two modes. A spatial permutation test (1,000 times) was applied to test the significance of spatial correlation. As shown in Figure 3A and Figure 3B, similar spatial correlations were observed between the loading pattern of the Association Development mode and distributions of cognitive-control- (e.g., “Cognitive-control & task performance”) and language-related (e.g., “Reading & Words”) brain activation connected by WM tracts. Furthermore, the loading patten of the Subcortical/limbic Development mode was correlated with reward or emotion related topics such as “Depression,” “Fear Conditioning & PTSD,” and “Reward” (Figure 3A, Figure 3B and Table S7).

**Figure 3.**
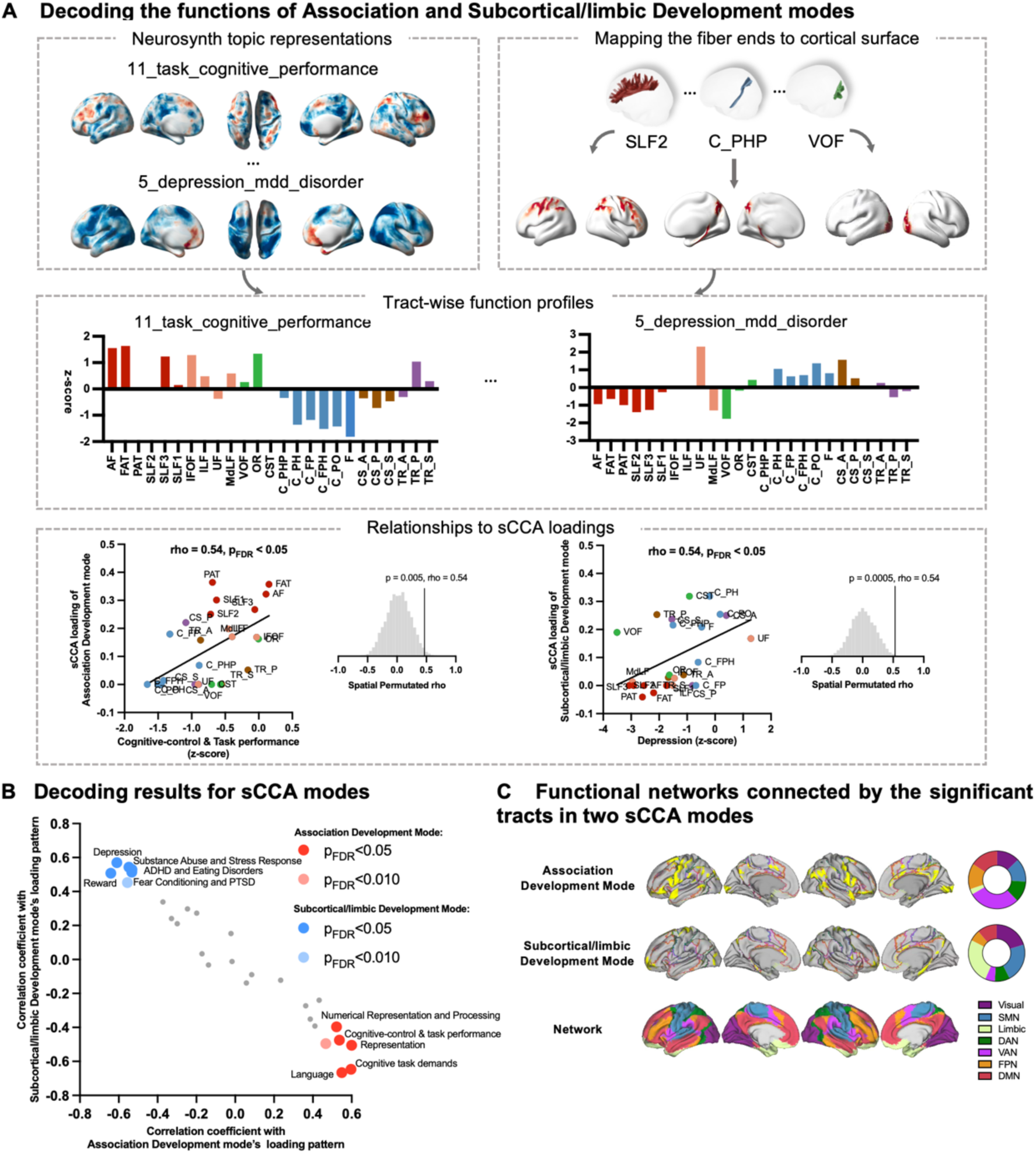
Results of tract-based functional decoding analysis. (A) Workflow for evaluating meta-analytic functional decoding of two sparse canonical correlation analysis (sCCA) loading patterns of the tract-based brain age gaps (BAGs). Selected Neurosynth topic maps represented on cortical surface were visualized in the left top panel. For each topic, a tract-wise functional profile was calculated by averaging z-scored brain functional activation probability within the gray matter (GM) regions connected by each white matter (WM) tract. In the right panel, representative scatterplots of spatial correlations between tract-wise functional profiles and sCCA loading patterns are shown. (B) Functional decoding results for two sCCA loading patterns. Spearman correlation coefficients with two sCCA loading patterns are plotted on x- and y-axes, respectively. The significantly correlated topics are highlighted and labeled. (C) Proportions of the functional networks connected by the WM tracts significant in the two sCCA modes. Pie plots demonstrate the percentages of tract-connected cortical voxels located in the seven resting-state cortical networks for the two sCCA modes. See Table S1 for the full names of the tracts. Abbreviations: SMN, sensorimotor network; DAN, dorsal attention network; VAN, ventral attention network; FP, frontoparietal network; DMN, default mode network; FDR, false discovery rate; PTSD, Posttraumatic Stress Disorder; ADHD, Attention-deficit/hyperactivity Disorder.

To qualitatively compare differences in functional networks connected by the tracts significantly loaded on the two sCCA modes, we projected their connected brain regions onto the cortical surface and overlapped with 7-network parcellation by Yeo et. al^34^. We observed that high-order networks (e.g., VAN, FPN and DMN) had the largest proportions of the brain regions connected by the tracts in the Association Development mode. Moreover, the tracts in the Subcortical/limbic Development mode connected mostly to the brain regions in sensorimotor and limbic networks (Figure 3C).

### Mitochondrial correlates of two cognition-psychopathology profiles

To explore potential biological mechanisms underlying the two cognition–psychopathology profiles, we incorporated six high-resolution ex-vivo mitochondrial maps, including 3 enzymatic activity maps (CI: NADH–ubiquinone oxidoreductase; CII: succinate dehydrogenase; CIV: cytochrome oxidase) reflecting energy-transformation, 1 mitochondrial content map (MitoD), and 2 derived maps representing tissue respiratory capacity (TRC; 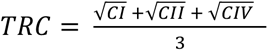) and mitochondrial respiratory capacity (MRC; MRC = TRC/MitoD)^25^ (see Methods). MRC refers to the ability of mitochondria to generate energy through oxidative phosphorylation, reflecting the energetic demands and metabolic support of specific tracts^25^. Tract-wise spatial mitochondrial distributions were quantified by overlaying the mitochondrial maps onto the *HCP-1065 tract atlas*^35^, from which mean mitochondrial values were extracted for each tract (Figure 4A). Statistical significance of each mitochondrial profile was assessed using a spatial spin-permutation null model (1,000 iterations). As shown in Figure 4B, WM tracts with higher loadings in the Association Development mode were significantly associated with increased enzymatic activity (CI: rho = 0.49, CII: rho = 0.47, CIV: rho = 0.52, *p_FDR_* < 0.05), elevated mitochondrial content (rho = 0.50, *p_FDR_* < 0.05) and higher MRC (rho = 0.48, *p_FDR_* < 0.05).

**Figure 4.**
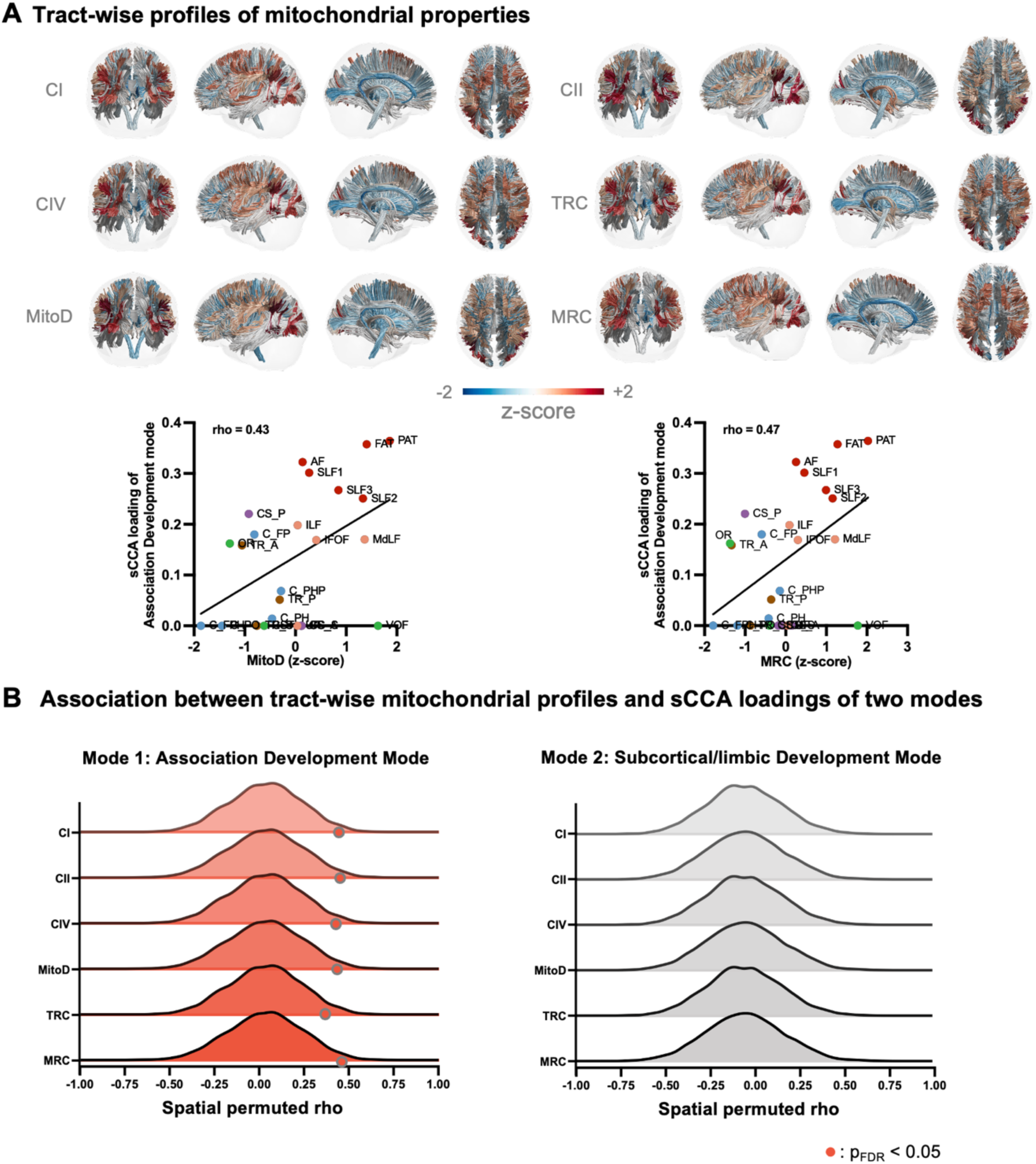
Results of tract-based mitochondrial decoding analysis. (A) Visualization of tract-wise mitochondrial profiles and spatial correlations with sparse canonical correlation analysis (sCCA) loadings for the Association Development Mode. (B) Association between tract-wise mitochondrial profiles and sCCA loadings of two modes. The distributions of rho values generated by spin-permutation tests are plotted. The mitochondrial map showing significant association with sCCA loading pattern is highlighted. See Table S1 for the full names of the tracts.

### Advanced maturation in WM tracts predicted cognitive performance two to three years later

Having established the associations between maturation of WM tracts and both cognitive function and psychopathology, we next investigated whether developmental deviations in WM tracts predict cognitive performance at follow-ups. To avoid circularity, we selected general cognitive measures^36^ that were not included in the prior sCCA analyses. Specifically, we included math ability (assessed via Stanford Mental Arithmetic Response Time Evaluation [SMARTE]), school grade and the overall performance of emotional Stroop task (measured by overall accuracy [STRP_ACC]), from a 2- or 3-year-follow-up (Figure 5A). For the relationships with these three follow-up cognitive measures, the Association Development mode demonstrated greater effect sizes (School Grade: 𝐹_(1, 7868)_ = 59.3, 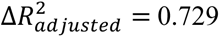, *p_FDR_* < 0.001; SMARTE: 𝐹_(1, 5623)_ = 102.9, 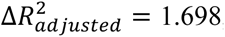, *p_FDR_* < 0.001; STRP_ACC: 𝐹_(1, 5967)_ = 33.6, 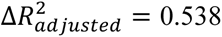, *p_FDR_* < 0.001) compared to those of Subcortical/limbic development mode (School Grade: 𝐹_(1, 7868)_ = 24.6, 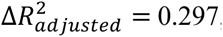, *p_FDR_* < 0.001; SMARTE: 𝐹_(1, 5623)_ = 69.6, 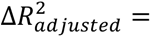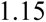, *p_FDR_* < 0.001; STRP_ACC: 𝐹_(1, 5967)_ = 14.3, 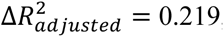, *p_FDR_*< 0.001). Tract-based BAGs with significant associations with the three cognitive measures are shown in Figure 5B and Table S8-10. Among the three cognitive performances, greater WM tract maturation was predictive of better cognitive performance assessed 2 to 3 years later.

**Figure 5.**
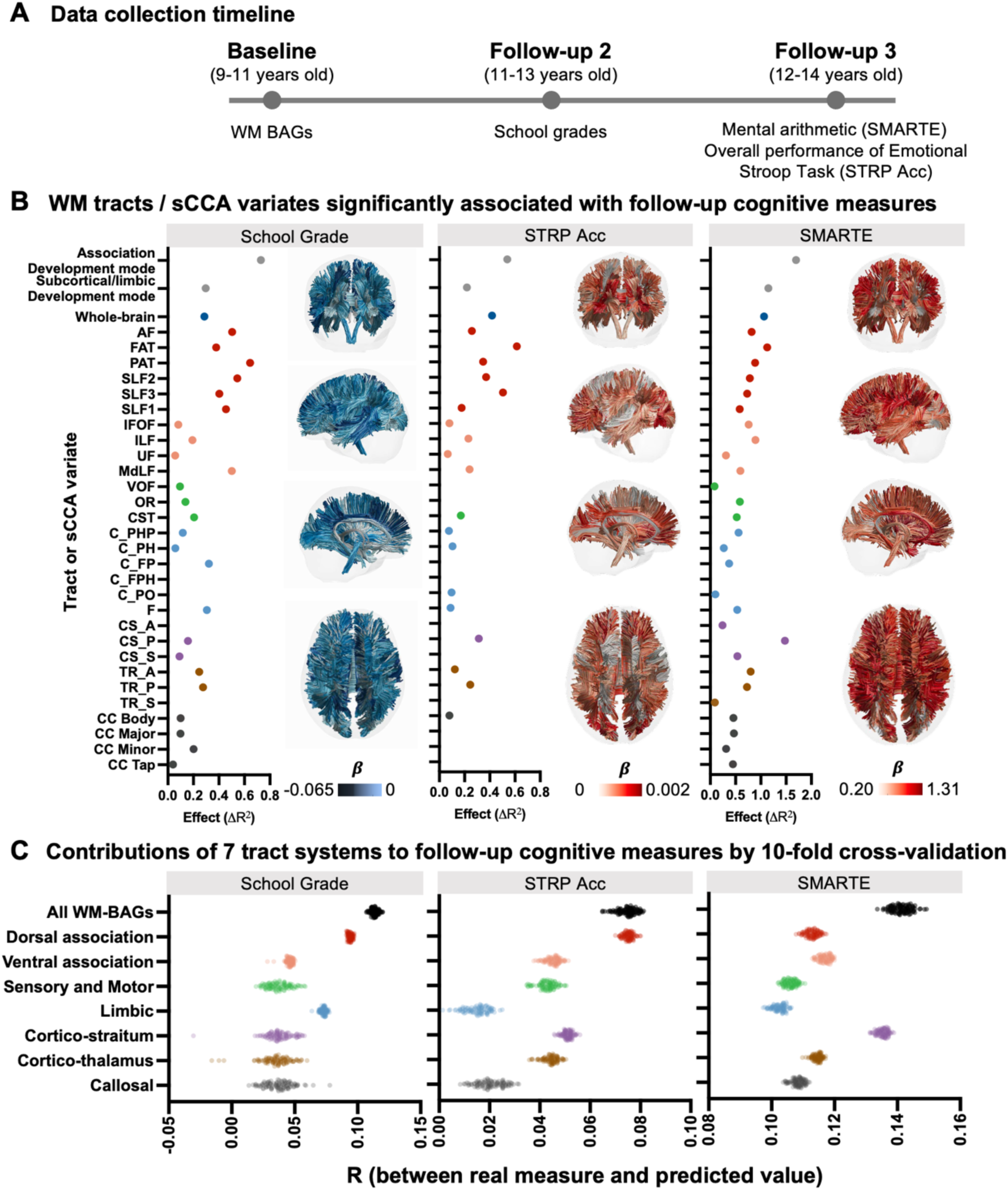
Relationships between baseline tract-BAG measures and cognitive performance at follow-ups. (A) Data collection timeline for cognitive measures at follow-ups. School grade was measured at 2-year follow-up, while inhibitory control and math ability were assessed at 3-year follow-up. (B) Relationships between tract-based measures and follow-up cognitive measures. Only the tract-based measures showing significant associations with cognitive measures are plotted. Effect sizes (Δ𝑅^2^) were calculated based on the change in overall proportion of variance by adding the predictor into the covariates-only model. The color of each tract on the right side of the sub-panel corresponds to the GAM coefficient of tract-based BAGs with significance. (C) Cross-validation results for different tract-based BAGs within eight systems. The scatter plots illustrate the distribution of correlation (R) values between the predicted and true cognitive scores over 100 iterations. See Table S1 for the full names of the tracts. Abbreviations: Stanford Mental Arithmetic Response Time Evaluation; STRP_Acc: emotional Stroop task accuracy; GAM: generalized additive model.

To further evaluate the contributions of tract-based BAGs in different systems to the prediction performance of follow-up cognitive measures, we built eight multivariate predictive models using the support vector regression (SVR) algorithm and a 10-fold cross-validation framework, repeated across 100 random iterations for each cognitive measure. Seven tract systems and the whole-brain tract system (with all 30 tract-based BAGs) were included. Model performance was assessed by Pearson’s correlation coefficient between real cognitive measures and the predicted cognitive measures during each iteration. As shown in Figure 5C, the whole-brain tract system had the highest predictive performance for both school grade and SMARTE scores across all eight systems. Among the individual tract systems, the dorsal association tracts provided the strongest predictive performance for school grades, while the cortico-striatal tracts were most predictive for SMARTE scores. For STRP_ACC, both the whole-brain and dorsal association systems demonstrated superior predictive performance compared to the other tract systems.

### Delayed maturation of WM tracts assessed via BAG at baseline concurrently associated with greater number of psychiatric (KSADS-5) diagnoses

We assessed whether tract-based BAGs were associated with psychiatric diagnoses using general linear models (GLMs). 8,594 participants with valid dMRI and psychiatric assessments from the ABCD dataset at baseline were used for the main analysis. For each participant, categorical psychiatric diagnoses were measured using the parent-reported Kiddie Schedule for Affective Disorders and Schizophrenia for DSM-5 (KSADS-5). sCCA variate scores derived from the Association Development mode at baseline showed a significant association with the cumulative numbers of KSADS-5 diagnoses assessed concurrently ( 𝐹_(2, 8591)_ = 15.06, *p_FDR_* < 0.001, Figure 6A-left), such that individuals, with less tract maturation relative to chronological age had higher number of psychiatric disorders. sCCA variate scores derived from the Subcortical/limbic Development mode showed a smaller effect for the association (𝐹_(2, 8591)_ = 6.22, *p_FDR_* = 0.008, Figure 6A-left). Moreover, we conducted analyses examining the temporal stability of these associations in 5,148 participants from ABCD dataset who had both tract-BAG and KSADS-5 diagnoses assessed at 2-year follow-up. Results of this analysis showed that individuals with lower BAGs (i.e., less tract maturation) from both modes were associated with more cumulative numbers of diagnoses at 2-y follow-up (Figure S5A and Table S11).

**Figure 6.**
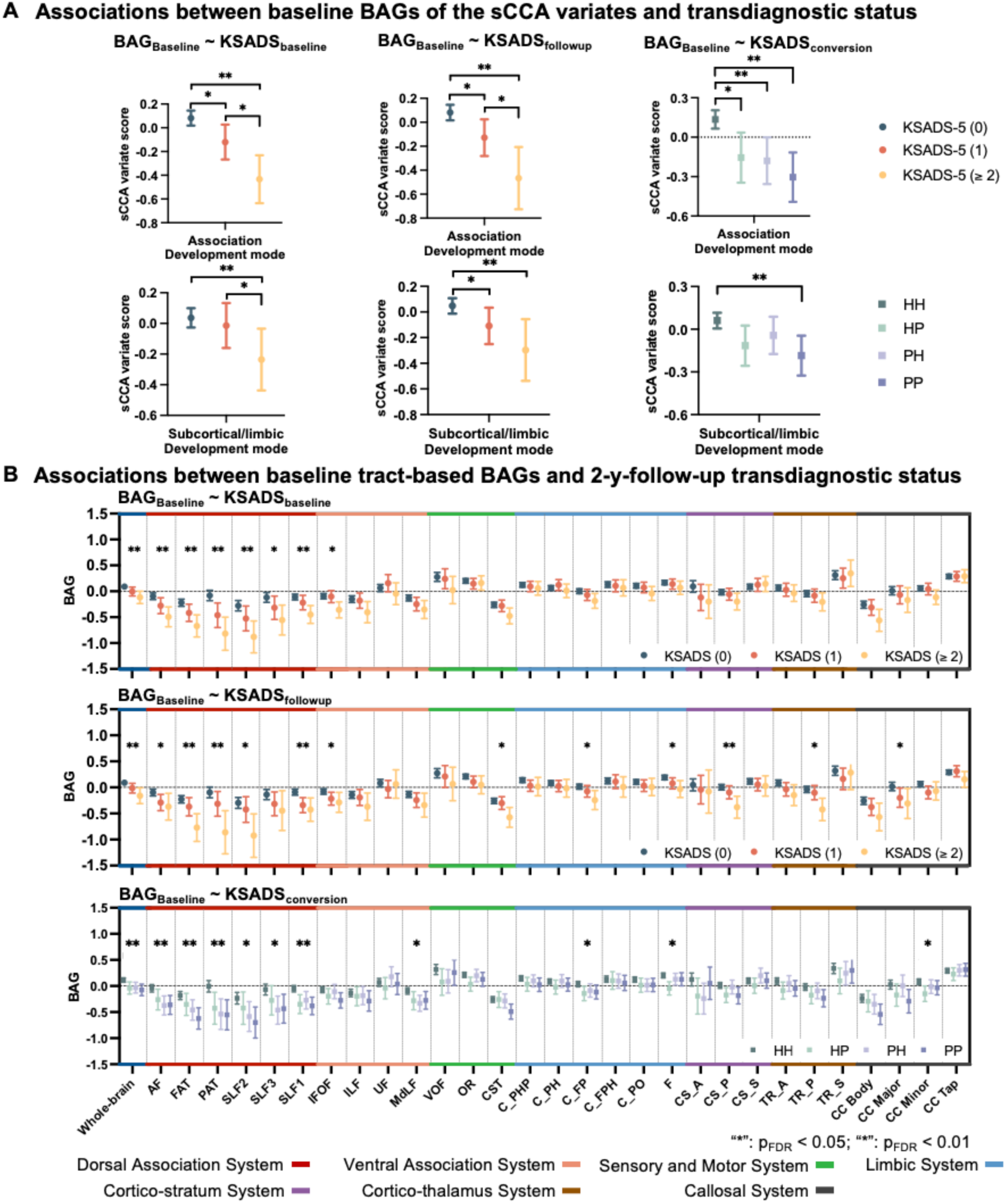
Relationships of baseline tract-BAG measures with the cumulative number of KSADS-5 diagnoses at baseline, at 2-y-follow-up, and the transition of the transdiagnostic status between baseline and 2-y-follow-up. Group-wise comparisons were conducted by generalized linear model (GLM) analyses. (A) Groupwise comparison of the baseline sparse canonical correlation analysis (sCCA) variate scores with the cumulative number of KSADS-5 diagnoses at baseline, 2-y-follow-up and 2-y-follow-up transdiagnostic status transitions. (B) Associations between baseline tract-based BAGs and the cumulative number of KSADS-5 diagnoses at baseline, 2-y-follow-up and 2-y-follow-up transdiagnostic status transitions. The top two panels show the tract-BAG sCCA variate scores of the groups with different cumulative number of KSADS-5 diagnoses at baseline and 2-y-follow-up. The bottom panel displays the tract-BAG sCCA variate scores of four groups including Healthy-persistent (HH), Disorder-remitted (PH), Disorder-new-onset (HP) and Disorder-persistent (PP) group. Error bars represent 95% confidence intervals. “*” *p_FDR_* < 0.05; “**” *p_FDR_* < 0.01. See Table S1 for abbreviations of the tracts. Abbreviations: BAG, brain-age gap; FDR, false discovery rate; KSADS-5: Kiddie Schedule for Affective Disorders and Schizophrenia for DSM-5 (KSADS-5).

In addition, we also investigated the relationships of BAGs obtained from individual tracts and whole-brain tracts, respectively, with the clinical diagnoses. BAG of the whole-brain tracts at baseline was significantly associated with the cumulative number of KSADS-5 diagnoses (0, 1, or >=2) assessed at the same time, after FDR correction (KSADS-5 effect: 𝐹_(2, 8591)_ = 7.05, *p_FDR_* = 0.004, Figure 6B-upper). Moreover, BAGs of all tracts within dorsal association system at baseline showed significant associations with the cumulative number of diagnoses assessed concurrently (e.g., FAT: 𝐹_(2, 8591)_ = 9.78, *p_FDR_* = 0.001 and PAT: 𝐹_(2, 8591)_ = 13.9, *p_FDR_* < 0.001, Figure 6B-upper). Post-hoc comparisons showed significantly more negative BAGs in participants with more diagnoses compared to those with no diagnosis. See Table S12 for results of other tracts and post-hoc analyses.

### Deviation in WM maturation assessed via BAGs at baseline prospectively predicted cumulative number of psychiatric (KSADS-5) diagnoses at 2-year follow-up

Next, the GLM analysis revealed that the scores derived from both sCCA variates at baseline were significantly associated with the cumulative number of diagnoses at 2-year follow-up (Association Development mode: 𝐹_(2, 7904)_ = 11.90, *p_FDR_* < 0.001; Subcortical/limbic Development mode: 𝐹_(2, 7904)_ = 7.20, *p_FDR_* < 0.001, Figure 6A-middle). Further, post-hoc analyses contrasting the difference between each pair of the subgroups showed that the scores of both sCCA variates were significantly lower in the subgroups with two or more diagnoses compared to the subgroup with no diagnosis.

BAG of the whole-brain tracts at baseline were significantly associated with the cumulative number of diagnoses at 2-year follow-up (𝐹_(2, 7904)_ = 7.03, *p_FDR_* = 0.005, Figure 6B-middle), with the post-hoc result of significantly more negative BAG in the subgroup with 2 or more diagnoses compared to the subgroup with no diagnosis. BAGs of 6 association tracts and 5 limbic/cortico-subcortical tracts at baseline showed significant associations with the cumulative number of diagnoses at 2-year follow-up (e.g. FAT: 𝐹_(2, 7904)_ = 8.64, *p_FDR_* = 0.002; CS_P: 𝐹_(2, 7904)_ = 7.91, *p_FDR_* = 0.003; Figure 6B-middle and Table S12). Post-hoc analyses revealed significantly more negative BAGs (i.e., less maturation) in the subgroup with 2 or more diagnoses compared to the subgroup with no diagnosis.

### Deviation in WM maturation assessed via BAGs at baseline prospectively predicted psychiatric diagnosis status transitions from baseline to 2-year follow-up

Brain variate scores of both sCCA modes at baseline significantly associated with the status transitions of - diagnoses from baseline to 2-year follow-up (Association Development mode: 𝐹_(3, 7826)_ = 10.52, *p_FDR_* < 0.001 ; Subcortical/limbic Development mode: 𝐹_(3,7826)_ = 4.75, *p_FDR_* = 0.008, Figure 6A-right). Post-hoc subgroup-wise comparisons showed that the brain variate scores of both sCCA modes were significantly lower in Disorder-persistent (Association Development mode: 𝑧 = −4.410 ; 𝑝 < 0.001 ; Subcortical/limbic Development mode: 𝑧 = −3.685; 𝑝 = 0.001) compared to Healthy-persistent subgroup, while the brain variate scores of the Association Development mode were significantly lower in Disorder-new-onset and Disorder-remitted subgroups compared to Healthy-persistent subgroup.

BAG of the whole-brain tracts at baseline showed a significant association with diagnostic status transitions (𝐹_(3, 7826)_ = 6.06, *p_FDR_* = 0.002, Figure 6B-bottom). For BAGs of individual tracts, all the tracts in the dorsal association system (e.g. AF: 𝐹_(3, 7826)_ = 7.04, *p_FDR_* = 0.001 ; FAT: 𝐹_(3, 7826)_ = 7.27, *p_FDR_* = 0.001 ) (Figure 6B-bottom and Figure S6) were significantly associated with diagnostic status transitions. See Table S12 for other tracts.

### Sensitivity analysis for the effects of puberty, socioeconomic status (SES) and age-bias correction

We conducted several sensitivity analyses to evaluate the robustness of our findings and determine if they remained significant after controlling for potential confounding factors. First, to test whether age-bias correction influenced the magnitude and significance of associations observed between WM BAGs and psychiatric diagnoses, we repeated the analyses using tract-based BAGs without age-bias correction. The results (Table S13) remained consistent, indicating that the observed associations were not dependent on the bias correction procedure.

Second, we sought to account for potential effects of pubertal development on our outcomes, by re-running our main analyses including Pubertal Development Scale (PDS) scores as an additional covariate. This analysis yielded unchanged results, suggesting that puberty did not confound the observed associations (Table S14).

Lastly, to account for effects related to SES on our outcomes, we reran our main analyses including two measures of SES (family income and parental education) as covariates. The results of this analysis (Table S15) were unchanged from our main results, indicating that the associations observed between tract-based BAG and psychiatric diagnoses could not be better accounted for by SES-related factors.

## Discussion

In this study, we investigated links of WM tract development during preadolescence with cognitive performance and transdiagnostic dimensions of psychological functioning. First, we established and validated normative models of development for specific and whole-brain WM tracts using three large, independent dMRI datasets. Critically, we are the first to show that chronological age during development can be significantly predicted using tract-based imaging features as well as whole-brain features. Next, we identified two distinct tract-BAG-based latent variates, weighted on the developmental levels of association tracts and subcortical/limbic tracts respectively, using a multivariate correlation analysis, that were associated with a wide range of cognitive and psychopathological measures. In both tract-based development modes, more mature WM tract systems were linked with greater cognitive function. In contrast, the maturation of association tracts was linked to lower psychopathology symptoms as measured by the CBCL, while the development of subcortical/limbic tracts was positively related to lower manic symptoms. Then, we showed that the maturation of association tracts was more predictive of follow-up cognitive performance than other individual tracts. Finally, we found that delayed maturation of the dorsal association tracts was concurrently and prospectively associated with the number of psychiatric diagnoses at baseline and 2-year follow-up, and predicted diagnostic status changes over that 2- year period. Collectively, the overlap in findings among these distinct analyses provides strong evidence that general cognitive performance and transdiagnostic psychiatric disorders are associated with WM maturation in the association tracts during preadolescence.

### Deviations in WM tract maturation assessed via BAG during preadolescence

Development trajectories of WM tracts have been extensively studied, indicating significant WM growth throughout childhood and adolescence^37, 38^. A novel contribution of our current study is that we demonstrated that chronological age was significantly predicted using tract-based dMRI features combined with machine learning algorithms. Our whole-brain model achieved an R^2^ of 0.70 and MAE of 1.81 years without age-bias correction in cross-validation. These results are comparable to other brain-age prediction studies in adolescents^27, 39, 40, 41^. Further, our study’s use of three independent datasets is a major strength reflecting the generalizability of the findings. We conducted a unique characterization on individual development of 29 WM tracts extracted from dMRI data with high reliability^35^. Using diffusion features of individual WM tracts, chronological age was significantly predicted in independent testing datasets. Interestingly, individuals with similar BAGs predicted by the whole-brain WM tracts showed varying tract-specific BAG profiles. Thus, our tract-based BAG approach is capable of providing unique insights into typical or atypical WM development and psychiatric risk.

### Tract-based BAGs define dimensions shared by cognition and psychopathology

Our sCCA analyses identified two brain-behavior modes (Association Development and Subcortical/limbic Development modes), linking advanced maturation of WM tracts with better cognitive function and less risk of psychopathology (Fig. 2). More specifically, greater development of WM association tracts was significantly linked with lower total psychological and attentional problems. More mature subcortical/limbic tracts are specifically associated with better decision-making and inhibitory control and lower manic symptoms. Post-hoc univariate correlation analysis also showed that individuals with advanced maturation of WM tracts generally had better cognitive performance and reduced psychiatric vulnerability (Fig. S3-4). Over the last decade, growing evidence has linked the development of WM microstructure to high-order cognitive abilities such as working memory, inhibition and cognitive shifting^42^. A study using the ABCD dataset also showed that a latent factor derived from executive functions was positively linked to higher FA in nearly all the involved WM tracts^43^, which fits with the wealth of findings linking WM deficits and various psychiatric disorders^19, 44^. Additionally, WM deficits were proposed to occur prior to illness onsets and remain stable with increasing age, suggesting that psychiatric disorders are neurodevelopmental in nature^45^. Our results align with a prior study showing in a sample of age-matched preadolescents^13^ that those with more maturation of WM tracts had more efficient communication among distinct brain regions, while those with delayed maturation of WM tracts were more likely exhibit signs/symptoms of psychiatric disorders.

This work also proposes a novel framework that integrates meta-analytic brain activation and mitochondrial/chemoarchitectural maps to elucidate neurobiological mechanisms of specific WM tracts. By analyzing the brain task-fMRI activations in regions connected by these tracts, we identified that the BAGs of Association Development mode are associated with cognitive-control, language and attention functions, while the Subcortical/limbic Development mode correlates with emotion processing and psychiatric symptoms. Our results showing cognitive and psychopathology associations with WM BAGs specific to association and subcortical/limbic tracts reinforces previous brain-behavior findings and offer new insights into how WM development contributes to distinct brain activation patterns. Previous studies have shown that functional connectivity of the association system (e.g., frontoparietal and salience-ventral attention networks) contributed more to higher-order cognition^32, 46^. In addition, abnormal (hypo- or hyper-) functional activation was evident particularly in the association system (e.g., salience-ventral attention networks) in transdiagnostic psychiatric disorders^6, 47^. Consistent with this model, our results demonstrated that greater development of association tracts captured the largest variations in the behavioral set and related more to higher-order cognition and lower total psychopathology. In addition, our tract-based mitochondrial decoding analysis identified mitochondrial enrichment in WM tracts from our sCCA modes implicated in cognition and psychopathology (Fig. 4). Specifically, we found that the WM tracts with high loadings in the Association Development Mode had higher mitochondrial content and energy metabolism than other WM tracts. Brain regions exhibiting higher MRC are known to have emerged later in evolutionary timeline^25^, suggesting increased vulnerability during neurodevelopment from an evolutionary psychiatry perspective^48^. WM tracts connecting association cortices undergo a more protracted development, reaching adult-level maturation at a later stage than those connecting sensory-motor cortices^11^. During early adolescence, WM tracts, particularly the association connections, have greater myelination, which requires enhanced mitochondrial activity and metabolism^9^. Consequently, WM tracts connecting association cortices may be particularly susceptible to mitochondrial dysfunction during this developmental window. Impaired mitochondrial function, as a result of exogenous or endogenous factors, may delay the maturation of these tracts, potentially compromising adolescents’ adaptive capacity in the face of cognitive and emotional challenges. Our findings align with post-mortem studies that have shown reduced WM mitochondrial density in patients with psychiatric disorders compared to non-psychiatric controls^14^, further supporting the “mitochondrial dysfunction” hypothesis in psychiatric disorders^15^.

Our findings revealed apparently contrasting associations between BAGs derived from WM and those derived from GM morphological features with respect to psychiatric symptom severity. Previous research has indicated that larger, positive BAGs based on GM features are often linked to more severe psychopathology, potentially driven by premature synaptic pruning^24^. It is noted that adolescence is within the second wave of dramatic myelin increase, which is essential for refining neural circuits to facilitate rapid information transmission^9^. A recent WM BAG study demonstrated that a more supportive educational environment accelerates the development of WM tracts crucial for academic skills^41^. Consistent with this, our study highlights the relationship between more advanced WM maturation, better cognitive performance, and reduced psychopathology. These findings suggest that WM BAG may serve as a neuroimaging marker of resilience, highlighting the differential roles of GM and WM development in mental health outcomes and cognitive function during adolescence.

### Maturation of specific tracts as a predictive biomarker of later cognitive performance

Our findings also highlight the potential of WM maturation in predicting cognitive performance two to three year later. Advanced WM maturation, particularly along the tracts implicated in the Association Development, was consistently associated with better cognitive performance. The superiority of this mode over Subcortical/limbic Development mode in predicting later-life cognitive performance suggests that more matured or optimized development of high-order association tract systems may have more long-term cognitive advantage. Multivariate predictive modeling further supports this point, revealing that using all tract-based BAG measures—and more specifically from dorsal association and cortico-striatal systems—provided the strongest predictive power for adolescents’ cognitive performance at follow-ups. The WM tracts within Association Development Mode connected the high-order brain networks of DMN, FPN and VAN. Integration of the triple networks including VAN, FPN and DMN networks is critical for maintaining cognitive-control ability in both adolescents and adults^49^. Cognitive-control ability includes inhibitory control, cognitive flexibility and working memory, which plays a vital role in high-order cognitive tasks like math performance, language and learning^50^. Previous functional study showed that higher functional connectivity levels of high-order brain networks, including DMN, FPN and VAN, were associated with better concurrent scholastic performance in participants aged 7 to 9 years old^51^. DTI studies in adolescents and adults also consistently found increased FAs in corticocortical and corticostiatal tracts were related to cognitive control ability^50, 52, 53^. Our findings verify the structural backbone of these high-order brain networks also be related to cognitive performance in adolescents and extend that the maturation level of structural connectivity at baseline have the predictive potential for follow-up cognitive performance.

### Maturation of specific tracts: insights into transdiagnostic and conversion status

In this study, we investigated the associations between tract-based BAGs and transdiagnostic status concurrently and at 2-year follow-up, as well as the conversion in diagnostic status, which reflects the change of transdiagnostic status across the 2-year follow-up period. We found that transdiagnostic dimensions of psychopathology and clinical status transitions from healthy to psychiatrically diagnosed states were both associated with delayed maturation of dorsal association tracts. The dorsal association tracts mainly connect the cortical brain areas of inferior parietal, frontal and superior parietal regions^35, 52^, largely overlapping with dorsal attention, ventral attention, and frontoparietal functional brain networks. This is consistent with our previous study showing that functional connectivity of the attention and frontoparietal systems is positively associated with cognition and negatively associated with psychopathology in preadolescents^32^. Moreover, reduced GM volume in the salience-ventral attention system is observed across individuals with psychiatric conditions in a disease non-specific manner, suggesting that disruptions in these networks have transdiagnostic implications^5, 54^. Conversely, in the current study, the WM tracts mainly connecting subcortical and limbic brain regions, which were related to reward-related measures, showed less predictive potential for the cumulative number of diagnoses concurrently and at 2-year follow-up. The development of limbic-, subcortical- and cortical-subcortical-based WM tracts (e.g. cingulum, fornix), involved in emotion regulation and decision-making, has shown a protracted developmental trajectory compared to projection and association tracts^11, 53, 55^. Their prolonged maturation period coincides with the onset of mood disorders^56^ and may suggest its critical role for supporting cognition-emotion integration.

### Limitations and future work

This study has several limitations. First, in this study, we primarily used data from subjects aged 8 to 11 years, with clinical and cognitive measures from the ABCD dataset at baseline. Given that there are different developmental windows for specific tracts^11^, it is plausible that links between psychopathology and developmental disruptions in other tracts would be evident if larger age ranges were assessed. Future studies should include further released follow-up data from the ABCD dataset and other developmental datasets with a broader age range to investigate how tract-specific developmental variability plays crucial roles in the neuropathology of transdiagnostic disorders at different life stages. Second, we limited the study to a subset of WM tracts with high robustness for auto extraction. Although we utilized the WM atlas with the most comprehensive definition of the tracts^35^, it did exclude some superficial WM tracts. More advanced fiber tracking and automatic tract extraction methods may improve the tract-based feature extraction for these superficial WM tracts. Third, our findings indicated that accelerated WM development was associated with better cognitive performance and lower transdiagnostic psychiatric dimensions, a relationship that contrasts with findings linking accelerated GM with greater psychiatric symptoms. Future multimodal BAG research is essential to examine how differing developmental trajectories of various neuroimaging markers such as WM and GM contribute to cognitive outcomes and psychiatric risk profiles. Fourth, our tract-wise mitochondrial profiles are currently based on a single postmortem adult brain that is not age/developmentally matched with our preadolescent sample. While it’s well known that mitochondrial content and function undergo changes across the lifespan^57, 58^, the utilized maps remain the only available postmortem dataset offering such high-resolution, voxel-wise mitochondrial data in WM^25^. To assess developmental trajectories of mitochondrial properties in WM, future work is needed to measure mitochondrial profiling in adolescent brains. In the present study, we primarily focused on the spatial patterning of tract-wise mitochondrial distribution, which is presumed to be relatively stable across age groups. Positron emission tomography studies have reported comparable spatial distributions of mitochondrial translocator protein (measured using [^11^C]PK11195) in both adults and children^59, 60^.

## Conclusion

In summary, our work showed that tract-specific WM features significantly predicted brain age for adolescents via a machine learning approach. We established tract-based BAGs, which enables deconvolution of the different developmental levels within an individual and measures brain development at tract-wise resolution. Using sCCA, we identified two distinct tract-BAG based latent brain variates, one loading on association tracts and the other on tracts connecting limbic or subcortical regions, which were associated with a wide range of cognitive and psychopathological measures. These findings provide early evidence that delayed maturation of WM tracts, especially in dorsal association tracts, may underly transdiagnostic risk for psychiatric disorders that emerge during preadolescence and early adolescence.

## Materials and Methods

### Participants

#### Human Connectome Project in Development (HCP-D) Study

The HCP-D dataset, designed to characterize healthy brain development in children and adolescents, was utilized as training and cross-validation for building a white-matter tract-based brain age model. This developmental dataset featured a predominantly cross-sectional design and comprised 652 individuals (aged 5.58 to 21.92 years; 351 females) with typical development^61^, of whom are without the following exclusion criteria: 1) Serious neurological condition; (2) History of serious head injury; (3) Long term use of immunosuppressants or steroids; (4) Premature birth; (5) Claustrophobia; (6) Hospitalization >2 days for certain physical or psychiatric conditions or substance use; (7) Treatment >12 months for psychiatric conditions or (9) Pregnancy or other MRI contraindications. All MRI data were acquired on 3T Siemens Prisma Scanners at four imaging sites. See more details about scanning parameters in Table S16 and Table S17. The study was approved by a central Institutional Review Board at Washington University in St. Louis.

#### Healthy Brain Network pediatric mental health (HBN) Study

To validate the prediction performance in an independent dataset with the similar age range, we included 1,687 subjects (600 females) from 5.58 to 21.90 years old from HBN study^62^. The MRI data was collected with 3T Siemens Prisma scanners at City College of New York and the CitiGroup Cornell Brain Imaging Center, and with a 3T Siemens Tim Trio scanner at Rutgers University Brain Imaging Center. See Table S16 and Table S17 for more scanning details. The study was approved by the Chesapeake Institutional Review Board.

#### Adolescent Brain Cognitive Development (ABCD) Study

We used the brain age model built from the HCP-D data to predict brain age for participants in the ABCD study, which is a prospective, ongoing, multi-site assessment of development in U.S. children from age 9-10 years though adulthood^63^. The MRI and behavioral data (v5.1) at two timepoints, baseline and 2-year-follow-up, were collected from 11,875 children. The T1-weighted (T1w) anatomical MRI and dMRI were acquired with magnetization-prepared rapid acquisition gradient echo (MPRAGE) and simultaneous multi-slice/multiband echo-planar imaging (EPI) sequences, respectively, using a unified protocol designed for different types of 3T MRI scanner platforms including Siemens Prisma and Prisma Fit, General Electronic Discovery 750, and Philips Medical System Ingenia and Achieva dStream. See Table S16 and Table S17 for more scanning parameter details.

### MRI data preprocessing and quality control (QC)

#### HCP-D

This dataset underwent the HCP’s minimal preprocessing workflow ^64^, including the steps of b0 intensity normalization, and corrections of EPI distortion, eddy-current,head-motion and gradient nonlinearity. The preprocessed files and QC metrics of 636 subjects were downloaded from the Fiber Data Hub (link: https://brain.labsolver.org/hcp_d.html). The data identified as “outlier” in the QC table were excluded (N =15), which were with lower neighboring diffusion weighted imaging (DWI) correlation values (0.46∼0.70).

#### HBN Study

The dMRI data was preprocessed by QSIPrep according to^65^. Preprocessed 3T dMRI data of 1,769 subjects were downloaded from AWS S3 (link: s3://fcp-indi/data/Projects/HBN/BIDS_curated/derivatives/qsiprep/). Because the dMRI data was collected according to 15 different acquisition parameters, we only included the subjects with the most common dMRI acquisition scheme of ‘SOTE_64dir_most_common’. Quality control (QC) procedure was from previous studies^65, 66^. The subjects whose dMRI quality score (’xgb_qc_score’, or ’dl_qc_score’ if ’xgb_qc_score’ was not available) less than 0.5 were excluded. A total of 1,102 subjects passed the quality control procedure.

#### ABCD Study

Standard dMRI preprocessing was performed by the ABCD Data Analysis, Informatics and Resource Center^67^, with the following steps of eddy-current correction, head-motion correction, adjusting diffusion gradients for head motion, robust tensor fitting, correcting B0 distortion and gradient distortion using opposite phase encoding pairs of b0 images, registering b0 images to T1w images using mutual information and cubic interpolation to resample at a 1.7 mm isotropic resolution. The QC procedure included both manual and automated checks, which were described elsewhere^67^. Specifically, we excluded the images that met one of the following criteria: (1) with artifacts preventing radiology reads or with incidental findings (mrif_score = 0, mrif_score = 3 or mrif_score = 4); (2) failed raw QC or total repetitions for all OK scans was less than 103 for dMRI data (iqc_dmri_ok_ser = 0 or iqc_dmri_ok_nreps < 103); (3) with dMRI B0 unwarp unavailable, with registration of dMRI to T1w larger than 17 mm, or dMRI maximum dorsal and ventral cutoff scores larger than 47 and 54 respectively (apqc_dmri_bounwarp_flag = 1, apqc_dmri_regt1_rigid > 17, apqc_dmri_fov_cutoff_dorsal > 47 or apqc_dmri_fov_cutoff_ventral > 54); (4) failed manual post-processing QC for dMRI data (dmri_dti_postqc_qc = 0) and (5) failed raw QC for T1w data (iqc_t1_ok_ser = 0). See Figure S7 for more details. In total, 8,773 participants at baseline and 5,494 at 2-year follow-up were used for the analyses.

### Assessments of cognition and psychopathology

The behavioral dataset, as used in our previous study^32^, comprised multisource data from the ABCD study assessing cognitive functioning and psychopathology across multiple domains. The dataset included 20 neurocognitive and 31 psychopathology-related assessments. The cognitive assessments covered different aspects of cognition including fluid cognition, crystalized cognition, overall cognition, processing speed, working memory, inhibitory control, episodic memory, vocabulary, reading, visuospatial ability, attention and cognitive flexibility, derived from NIH Toolbox cognition battery, Matrix Reasoning Test from the Wechsler Intelligence Scale for Children-V (WISC-V), Rey Auditory Verbal Learning Test (RAVLT), Little Man Task, Recognition memory, Emotional n-back fMRI task, Stop-signal fMRI task and Monetary Incentive Delay task. See Table S18 for more details of the above neurocognitive measurements. The psychopathology-related assessments included the scales of the Child Behavioral Checklist (CBCL)^68^, the personality measures from the Modified Urgency, Perseverance, Premeditation and Sensation-seeking (UPPS-P) for Children from PhenX^69, 70^, the mania scale from the 73-item Parent General Behavior Inventory (P-GBI), psychosis risk symptom subscales from Youth-report Prodromal Questionnaire Brief Version (PQ-B)^71^ and the subscales assessing two motivational systems from a modified Behavioral Inhibition & Behavioral Activation Scales. See Table S19 for details of psychopathology-related subscales.

### Assessments of clinical diagnosis and status transitions

We used parent-reported Kiddie Schedule for Affective Disorders and Schizophrenia for DSM-5 (KSADS-5)^72^ to measure the current categorical psychiatric diagnoses at baseline and 2-year follow-up. Psychiatric disorders (Depressive Disorder [MDD], Psychotic Disorder [PSD], Bipolar Disorder [BPD], Attention-deficit/hyperactivity Disorder [ADHD], Substance Use Disorder [SUD], Alcohol Use Disorder [AUD], Obsessive-compulsive Disorder [OCD], Social Anxiety Disorder [SAD], Generalized Anxiety Disorder [GAD], Separation Anxiety Disorder [SED], Eating Disorder [ED], Conduct Disorder [CD], Posttraumatic Stress Disorder [PTSD], Oppositional Defiant Disorder [ODD], Specific Phobia [SP], Panic Disorder [PAD], Agoraphobia [AGO], Disruptive mood dysregulation disorder [DMDD], Enuresis and Encopresis Disorder [EED], and Tic Disorder [TD]) were used for the analyses. For each specific disorder, the score would be labeled as “0” for “absence of diagnosis” and as “1” for “definitive diagnosis”.

### Diffusion MRI preprocessing

The dMRI data were reconstructed using generalized q-sampling imaging^73^ with a diffusion sampling length ratio of 1.25. A deterministic fiber tracking algorithm^74^ was used with augmented tracking strategies^75^ to improve reproducibility. We utilized default parameters for fiber tracking ^35^. Topology-informed pruning^76^ was applied to tractography with 2 iteration(s) to remove false connections. The above preprocessing was performed using DSI studio (https://dsi-studio.labsolver.org; ^35^).

### Tract profile quantification and data harmonization

A total of 54 tracts were automatically extracted using DSI studio. These tracts included 6 pairs of bilateral dorsal association tracts (left and right Arcuate Fasciculus [AF], Frontal Aslant Tract [FAT], Parietal Aslant Tract [PAT], Superior Longitudinal Fasciculus 1 [SLF1], Superior Longitudinal Fasciculus 2 [SLF2], Superior Longitudinal Fasciculus 3 [SLF3] tracts), 4 pairs of bilateral ventral association tracts (left and right Inferior Fronto Occipital Fasciculus [IFOF], Inferior Longitudinal Fasciculus [ILF], Uncinate Fasciculus [UF] and Middle Longitudinal Fasciculus [MdLF] tracts), 6 pairs of bilateral limbic tracts (left and right Cingulum Frontal Parahippocampal [C_FPH], Cingulum Frontal Parietal [C_FP], Cingulum Parahippocampal [C_PH], Cingulum Parahippocampal Parietal [C_PHP], Cingulum Parolfactory [C_PO] and Fornix [F] tracts), 3 pairs of bilateral sensory-motor tracts (left and right Corticospinal Tract [CST], Optic Radiation [OR] and Vertical Occipital Fasciculus [VOF] tracts), 3 pairs of bilateral thalamic radiation tracts (left and right Thalamic Radiation Anterior [TR_A], Thalamic Radiation Posterior [TR_P] and Thalamic Radiation Superior [TR_S] tracts), 3 pairs bilateral corticostriatal tracts (left and right Corticostriatal Tract Anterior [CS_A], Corticostriatal Tract Posterior [CS_P], and Corticostriatal Tract Superior [CS_S]) and 4 corpus callosum tracts (Corpus Callosum Body [CC_Body], Corpus Callosum Forceps Major [CC_Major], Corpus Callosum Forceps Minor [CC_Minor] and Corpus Callosum Tapetum [CC_Tap]). The 54 tracts are illustrated in Figure 1A. The GM regions mainly connected by the tracts are indicated in Table S1. For each identified tract, 100 equal-length segments (or nodes thereafter) were sampled along the tract from one end to the other (Figure 1B). We then computed the FA value for each node in all the tracts using a kernel density estimator with the default bandwidth setting. The 100-node FA tract-profile was generated for each tract of each participant. 611 participants in HCP-D dataset (aged 5.58 to 21.92 years; 281 females and 330 males), 978 participants in HBN dataset (aged from 5.58 to 21.90 years, 366 females), and 8,688 at baseline and 5,883 at 2-year follow-up in ABCD dataset (N_Baseline_ = 8,688, age range 8-11 years; N_2-year-follow-up_ = 5,883) had successful automatic tract extraction for all 54 WM tracts.

In order to eliminate the influence of the differences across acquisition sites, we harmonized the FA tract-profile data using a ComBat algorithm in Matlab (https://github.com/Jfortin1/ComBatHarmonization/tree/master/Matlab)^77^. In our study, KSADS-5 diagnosis, age and sex were designed as the biological variables to reserve.

### Brain age prediction models and brain age gap

#### Brain age prediction models

An overview of the brain age prediction pipeline is outlined in the Figure 1C. We used the HCP-D dataset for training and cross-validation of brain age prediction models, and then applied these models to predict brain age in the independent ABCD and HBN datasets. The 25 pairs of projection/association tracts (each pair of bilateral tracts were concatenated) and 4 corpus callosum tracts were used to build “tract-specific” models respectively, while all 54 tracts were concatenated to build a “whole-brain” model. Using a 5-fold cross-validation on 80% subjects from HCP-D dataset, a Gaussian Process Regression (GPR) model^26, 27^ (https://github.com/garedaba/brainAges) in Python was trained to predict individual age based on the whole-brain tract-profile or 29 tract-specific profiles respectively. The remaining 20% HCP-D subjects were used to validate the brain age model performance in a 5-fold cross-validation procedure.

The trained brain age prediction models from the HCP-D cohort were then used to predict brain ages of the subjects at baseline and 2-year-follow-up in the ABCD dataset and the subjects in the HBN dataset. Model accuracy was evaluated in the cross-validation dataset and the independent testing dataset using the R^2^ (the portion of the variance explained) and the MAE (mean absolute error).

#### Brain age gap calculation and brain-age bias correction

For each subject, brain age gap (BAG) was calculated as the difference between predicted brain age and chronological age. Age-bias correction was conducted by the method proposed by Beheshti et al.^29^. In the training dataset, we performed linear regression between chronological age and BAG, as follows:

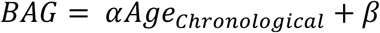

The derived slope (𝛼) and intercept (𝛽) were applied to the chronological ages of the individuals in the cross-validation and the independent testing datasets. We then subtracted the calculated values from the predicted BAGs to correct for age bias, as follows: 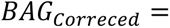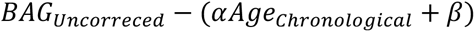

To avoid potential prediction accuracy overestimates^31^, we only reported the prediction performance by uncorrected brain ages. Tract-based BAGs without age-bias correction were used for testing the robustness of our results described in the statistical analysis section.

### Statistical analysis

#### Associations between tract BAGs and behavioral assessments

To explore the multivariate associations between the tract BAGs and the multidomain behavioral assessments, we performed sparse canonical correlation analysis (sCCA) between Z-scored brain and behavioral measurements of the ABCD baseline datasets (https://github.com/cedricx/sCCA/tree/master/sCCA/code/final)^78^. Prior to sCCA analysis, we regressed age and sex out of both the brain and behavioral measures. The sCCA workflow is shown in Figure 3A. sCCA identifies multiple latent variables that show maximal correlations between the brain and behavior datasets, with regularization to handle multicollinearity issues and achieve sparsity.

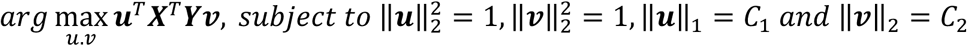

Here, X and Y are the multi-dimensional tract-based BAG and behavioral datasets, respectively. In addition, u and v are the coefficients used to project the datasets to latent variables. The regularization parameters include the sparsity of 𝒖 and 𝒗, which were determined by the following grid-search and nested cross-validation procedure within the training dataset. Both parameters were incremented by 0.1 from 0 to 1, resulting 100 combinations of different 𝐶_1_ and 𝐶_2_ values. Subsequently, we randomly resampled two-thirds of the subjects for ten times. Through two-thirds of the sample, the coefficients of the sCCA model with 𝐶_1_and 𝐶_2_were estimated. The combination of the regularization parameters with the largest canonical correlation coefficient was used for the final model. Furthermore, we performed 1,000 times of permutation tests to assess the statistical significance of canonical correlation modes, with shuffling the behavioral set order during each iteration. To determine the significance of each behavioral measure within each mode, we used a bootstrap resampling procedure, repeated 1,000 times. Behavioral measures for each sCCA mode were deemed significant if their 95% confidence intervals did not include zero.

Additionally, to assess the specificity of the sCCA modes, we performed paired t-tests to compare the behavioral loading distributions derived from the bootstrap tests. A behavioral measure was considered specific to an sCCA mode if its effect size (Cohen’s d value) was more than 0.5.

In addition, a post-hoc analysis investigating the relationships between tract BAGs and each cognitive or clinical score was conducted by generalized linear model (GLM) with the covariates of sex and age as follows:

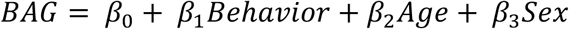

All tested associations (30 × 51 tests) were corrected for multiple comparisons by using the FDR method. Corrected p and t values are shown in Figure S3-4.

#### Associations between tract-BAG measures and clinical diagnoses at baseline

To assess clinical utility of the tract-BAG measures in the diagnoses of psychiatric disorders, we characterized associations of these BAGs with the cumulative number of KSADS-5 diagnoses in the ABCD dataset. We categorized the participants into three subgroups based on their cumulative number of their KSADS-5 diagnoses at baseline. Participants in the first subgroup were with no KSADS-5 diagnosis (N = 6,466), in the second subgroup with a single KSADS-5 diagnosis (N = 1,337), and in the third subgroup with at least two KSADS-5 diagnosis (N = 791). Tract-BAG measures (including sCCA variate scores and individual tract BAGs) versus cumulative number of KSADS-5 diagnoses were assessed using general linear models (GLMs) controlled for age and sex, as follows:

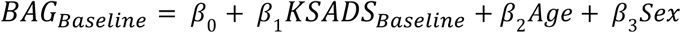

where categorical variable *KSADS_Baseline_* is the cumulative number of KSADS-5 diagnoses (0, 1, or ≥ 2) at baseline. We then performed Tukey tests for post-hoc comparisons. The GLM analysis was conducted using ‘glm’ in R.

We also conducted a validation analysis using GLMs to evaluate the associations between tract- BAG measures and transdiagnostic status in participants with available 2-year follow-up data. These participants, who had both brain and clinical measures available at the 2-year follow-up, were categorized into three groups: those without a KSADS-5 diagnosis (N = 4,131), those with one KSADS-5 diagnosis (N = 741), and those with at least two KSADS-5 diagnoses (N = 336).

#### Associations between tract-BAG measures at baseline and clinical diagnoses at 2-year follow-up

To explore the predictive potential of tract-BAG measures for clinical diagnoses of psychiatric disorders at 2-year-followup, we used GLMs to assess associations between the tract-BAG measures at baseline and cumulative number of KSADS-5 diagnoses at 2-year-followup, as follows:

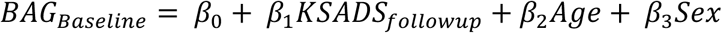

where variable *KSADS_followup_* is the cumulative number of KSADS-5 diagnoses at 2-year follow-up. The participants were categorized into three subgroups according: those without KSADS-5 diagnosis (N = 6,288), those with a single KSADS-5 diagnosis (N = 1,116) and those with at least two KSADS-5 diagnoses (N = 503).

#### Associations between tract-BAG measures at baseline and status transitions in KSADS-5 diagnoses from baseline to 2-year follow-up

We investigated whether tract-BAG measures associated with diagnostic status transitions. Based on whether the participants had psychiatric diagnoses at baseline and/or 2-year follow-up, they were categorized into 4 subgroups: Healthy-persistent (0 psychiatric diagnosis at both baseline and 2-year follow-up, *N*_𝐻𝐻_ = 5,236), Disorder-remitted (≥1 psychiatric diagnosis at baseline and 0 psychiatric diagnosis at 2-year follow-up, *N*_𝑃𝐻_ = 995 ), Disorder-new-onset (0 psychiatric diagnosis at baseline and ≥1 psychiatric diagnosis at 2-year follow-up, *N*_𝐻𝑃_ = 751) and Disorder-persistent (≥1 psychiatric diagnosis at both baseline and 2-year follow-up, *N*_𝑃𝑃_ = 848). GLMs were conducted to assess the association between tract BAGs versus the diagnostic transition status of psychiatric diagnosis, as follows:

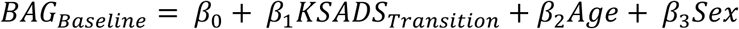

#### Associations between tract-BAG measures at baseline and cognitive performance at 2-year or 3- year follow-up

To evaluate longitudinal cognitive outcomes, we examined the associations between baseline tract-based BAG measures and follow-up cognitive performance not included in the prior sCCA analysis. Three general cognitive assessments from follow-up visits were used: Emotional Stroop Task (3-Year Follow-Up)^79^: This task assessed processing speed/attention, inhibitory control, and emotional information processing. We used the overall accuracy score as the measure. Self-Reported Academic Grade (SAG; 2-Year Follow-Up)^36^: School performance was evaluated using self-reported grades, ranging from 1 (A+) to 12 (failing), with lower scores indicating better academic performance. Stanford Mental Arithmetic Response Time Evaluation (SMARTE)^36^: This measure of math ability included dot enumeration, single-digit, and mixed-digit arithmetic problems. The total number of correct responses was used as the performance measure.

To assess the relationship between baseline tract-based BAGs and each follow-up cognitive measure, we employed generalized additive models (GAMs), which are widely used for modeling non-linear effects in large-scale datasets. Age and sex were included as covariates in all models. The unique contribution of each tract-based BAG measure was quantified by computing the change in model variance (adjusted ΔR², expressed as a percentage) when the BAG predictor was added to a null model containing covariates only. Both effect sizes and p-values were reported.

To further evaluate the predictive capacity of tract-based BAGs for cognitive performance, we applied support vector regression (SVR) within a 10-fold cross-validation framework, repeated 100 times for robustness. Eight combinations of tract-based BAG feature sets were evaluated: whole-brain (30 BAGs), dorsal association (AF, FAT, PAT, SLF2, SLF3 and SLF1 BAGs), ventral association (IFOF, ILF and UF BAGs), sensory and motor (CST, OR and VOF BAGs), limbic (C_PHP, C_PH, C_FP, C_FPH, C_PO and F BAGs), cortico-striatum (CS_A, CS_P and CS_S BAGs), cortico-thalamus (TR_A, TR_P and TR_S BAGs) and callosum (CC Body, Major, Minor and Tap BAGs). During each iteration, the data were randomly divided into a ‘training’ set (90% of subjects) and a ‘testing’ set (10% of subjects). The SVR model was trained on the ‘training’ set, and performance was evaluated on the ‘testing’ set. Prediction accuracy was measured by correlating true and predicted cognitive scores using Pearson correlation analysis.

#### Sensitivity analysis on effects of pubertal maturation

Pubertal maturation of the ABCD subjects was measured by the Pubertal Development Scale (PDS)^80^. We utilized the parent-reported PDS as it is likely more reliable than the self-reported version for the age range of our study participants and widely used in other ABCD studies^81^. Of 8,681 participants, 8,263 had pubertal development data assessed by the Pubertal Development Scale (PDS), which included puberty category measures ranging from prepubertal to post-pubertal. For male participants, the puberty category score was calculated by summing items related to voice deepening, body hair growth, and facial hair growth. These scores were then categorized as follows^82^: prepubertal (= 3), early pubertal (4 or 5), mid pubertal (≥ 6 and ≤ 8), late puberty (≥ 9 and ≤ 11); postpubertal (= 12). For female participants, the puberty category score was derived by summing items on body hair growth and breast development, with categorization as follows: prepubertal (= 2), early pubertal (=3 and no menarche), mid pubertal (> 3 and no menarche), late puberty (≤ 7 and menarche); postpubertal (= 8 and menarche). To assess the impact of pubertal maturation on our results, we included the puberty category as an additional covariate in the GLM analysis described above.

### Functional decoding based on spatial association between brain regions connected by WM tracts and task-related brain activation

To validate the functional significance of the tract-based BAGs of the sCCA modes, we assessed spatial correlation between brain regions connected by these tracts and specific task-related brain activation patterns. For task-related brain activation maps, we used data from the latest NeuroSynth database (version 7)^33^, a meta-analytic tool that aggregates findings from 14,371 published fMRI studies. We selected 31 topics (See Table S6) related to cognitive-control, language, emotion, or psychiatric conditions from a set of 100 topics generated using Latent Dirichlet Allocation (LDA). These topic maps were generated through reverse inference, denoting the probability of a topic being relevant given a particular brain activation. For the white matter tracts, we utilized the Population-Probability Atlas^35^, which provides voxel probability maps for 25 association and projection white matter tracts included in our sCCA study. To refine this atlas, we excluded voxels with a probability lower than 0.5 and those with less than 50% probability according to the GM probability map (*MNI-maxprob-thr50-2mm.nii.gz* in FSL). We calculated the average probability of topic activation levels within the generated tract regions of interest (ROIs). To evaluate their spatial correlation, we conducted Spearman correlation analyses between the loadings of two sCCA modes and the z-scores from the brain activation meta-analytic maps for each topic. Statistical significance was evaluated using a spin-based spatial permutation test (1,000 iterations; https://github.com/murraylab/brainsmash/tree/master)^83^, which preserves the spatial autocorrelation structure of GM maps. During each iteration, the voxel-wise task activation map was spatially rotated and the brain activation levels of the GM regions connected by each tract were recomputed. The empirical p-value was defined as the proportion of permuted Spearman correlation coefficients that exceeded the observed correlation and corrected by the FDR method.

### Mitochondrial mapping analyses

We based our analysis on whole-brain maps of mitochondrial properties developed by Mosharov et al.^25^. Six mitochondrial maps (CI, CII, CIV, MitoD, TRC and MRC) were included in our study. For each tract, we averaged the mitochondrial values within the voxels whose probability higher than 50% in Population-Probability Atlas^35^ in MNI152 volumetric space. The tract-wise mitochondrial profiles were shown in Figure 4A. Spatial correlation between sCCA loading pattern and mitochondrial profiles were performed by Spearman correlation. Statistical significance was evaluated using a spin-based spatial permutation test (1,000 iterations; https://github.com/frantisekvasa/rotate_parcellation)^84^, which preserves the spatial autocorrelation structure of the locations of the tracts. We then estimated the number of null correlation coefficients that were significantly higher than expected by chance as p-value and corrected by the FDR method.

## Acknowledge

This work was supported by the Intramural Research Program of the National Institute on Drug Abuse, National Institutes of Health (NIH) and utilized the computational resources of the NIH HPC Biowulf cluster (https://hpc.nih.gov).

The authors used data from the Adolescent Brain Cognitive Development Study (ABCD, abcdstudy.org), the Healthy Brain Networks (HBN, data.healthybrainnetwork.org) and he Lifespan Human Connectome Project Development Study (HCP-D, https://www.humanconnectome.org/study/hcp-lifespan-development). The ABCD Study, held in the National Institute of Mental Health (NIMH) Data Archive (NDA), is a multisite, longitudinal study designed to recruit more than 10,000 children aged 9–10 and follow them over 10 years into early adulthood. The ABCD Study is supported by the NIH and additional federal partners under award numbers U01DA041048, U01DA050989, U01DA051016, U01DA041022, U01DA051018, U01DA051037, U01DA050987, U01DA04 1174, U01DA041106, U01DA041117, U01DA041028, U01DA041134, U01DA050988, U01DA051039, U01DA041156, U01DA041025, U01DA 041120, U01DA051038, U01DA041148, U01DA041093, U01DA041089, U24DA041123, and U24DA041147. A full list of supporters is available at https://abcdstudy.org/federal-partners.html. A listing of participating sites and a complete listing of the study investigators can be found at https://abcdstudy.org/consortium_members/. HCP-D was supported by the NIMH of the NIH under Award Number U01MH109589 and by funds provided by the McDonnell Center for Systems Neuroscience at Washington University in St. Louis. The HCP-Development 2.0 Release data used in this report came from DOI: <u>10.15154/1520708</u>. Preprocessed SRC files of HCP-D dataset were downloaded from Fiber Data Hub (https://brain.labsolver.org/hcp_d.html). HCP-D, HBN and ABCD consortium investigators designed and implemented the study and/or provided data but did not necessarily participate in analysis or the writing of this report. This manuscript reflects the views of the authors and may not reflect the opinions or views of any other agency, organization, employer or company.

## Competing Interest Statement

The authors declare that they have no known competing financial interests or personal relationships that could have appeared to influence the work reported in this paper.

## Notes

### Competing Interest Statement

The authors have declared no competing interest.

## Reference

1. Substance Abuse and Mental Health Services Administration. Key substance use and mental health indicators in the United States: Results from the 2021 National Survey on Drug Use and Health. HHS Publication No PEP22-07-01-005, NSDUH Series H-57, (2022).

2. Walker ER, McGee RE, Druss BG. Mortality in mental disorders and global disease burden implications: a systematic review and meta-analysis. JAMA Psychiatry 72, 334–341 (2015).

3. Hyman SE. The daunting polygenicity of mental illness: making a new map. Philos Trans R Soc Lond B Biol Sci 373, (2018).

4. Lee SH, et al. Genetic relationship between five psychiatric disorders estimated from genome-wide SNPs. Nature Genetics 45, 984-+ (2013).

5. Goodkind M, et al. Identification of a Common Neurobiological Substrate for Mental Illness. JAMA Psychiatry 72, (2015).

6. McTeague LM, Huemer J, Carreon DM, Jiang Y, Eickhoff SB, Etkin A. Identification of Common Neural Circuit Disruptions in Cognitive Control Across Psychiatric Disorders. American Journal of Psychiatry 174, 676–685 (2017).

7. Solmi M, et al. Age at onset of mental disorders worldwide: large-scale meta-analysis of 192 epidemiological studies. Molecular Psychiatry 27, 281–295 (2021).

8. Paus T, Keshavan M, Giedd JN. Why do many psychiatric disorders emerge during adolescence? Nature Reviews Neuroscience 9, 947–957 (2008).

9. de Faria O, Pivonkova H, Varga B, Timmler S, Evans KA, Káradóttir RT. Periods of synchronized myelin changes shape brain function and plasticity. Nature Neuroscience 24, 1508–1521 (2021).

10. Riccomagno MM, Kolodkin AL. Sculpting neural circuits by axon and dendrite pruning. Annu Rev Cell Dev Biol 31, 779–805 (2015).

11. Sydnor VJ, et al. Neurodevelopment of the association cortices: Patterns, mechanisms, and implications for psychopathology. Neuron 109, 2820–2846 (2021).

12. Kochunov P, et al. White Matter in Schizophrenia Treatment Resistance. Am J Psychiatry 176, 829–838 (2019).

13. Karlsgodt KH. White Matter Microstructure across the Psychosis Spectrum. Trends Neurosci 43, 406–416 (2020).

14. Senko D, et al. White matter lipidome alterations in the schizophrenia brain. Schizophrenia (Heidelb) 10, 123 (2024).

15. Manji H, et al. Impaired mitochondrial function in psychiatric disorders. Nat Rev Neurosci 13, 293–307 (2012).

16. Kelly A, et al. Age-Based Reference Ranges for Annual Height Velocity in US Children. The Journal of Clinical Endocrinology & Metabolism 99, 2104–2112 (2014).

17. Halperin JM, McKay KE. Psychological Testing for Child and Adolescent Psychiatrists: A Review of the Past 10 Years. Journal of the American Academy of Child & Adolescent Psychiatry 37, 575–584 (1998).

18. Marquand AF, Kia SM, Zabihi M, Wolfers T, Buitelaar JK, Beckmann CF. Conceptualizing mental disorders as deviations from normative functioning. Molecular Psychiatry 24, 1415–1424 (2019).

19. Tung YH, et al. Whole Brain White Matter Tract Deviation and Idiosyncrasy From Normative Development in Autism and ADHD and Unaffected Siblings Link With Dimensions of Psychopathology and Cognition. Am J Psychiatry 178, 730–743 (2021).

20. Chien Y-L, et al. Neurodevelopmental model of schizophrenia revisited: similarity in individual deviation and idiosyncrasy from the normative model of whole-brain white matter tracts and shared brain-cognition covariation with ADHD and ASD. Molecular Psychiatry 27, 3262–3271 (2022).

21. Bethlehem RAI, et al. Brain charts for the human lifespan. Nature 604, 525–533 (2022).

22. Kaufmann T, et al. Common brain disorders are associated with heritable patterns of apparent aging of the brain. Nat Neurosci 22, 1617–1623 (2019).

23. Schnack HG, van Haren NE, Nieuwenhuis M, Hulshoff Pol HE, Cahn W, Kahn RS. Accelerated Brain Aging in Schizophrenia: A Longitudinal Pattern Recognition Study. Am J Psychiatry 173, 607–616 (2016).

24. Cropley VL, et al. Brain-Predicted Age Associates With Psychopathology Dimensions in Youths. Biological Psychiatry: Cognitive Neuroscience and Neuroimaging 6, 410–419 (2021).

25. Mosharov EV, et al. A human brain map of mitochondrial respiratory capacity and diversity. Nature 641, 749–758 (2025).

26. Rasmussen CE. Gaussian processes in machine learning. Lect Notes Artif Int 3176, 63–71 (2004).

27. Ball G, Kelly CE, Beare R, Seal ML. Individual variation underlying brain age estimates in typical development. NeuroImage 235, (2021).

28. Cole JH, et al. Brain age predicts mortality. Molecular Psychiatry 23, 1385–1392 (2017).

29. Beheshti I, Nugent S, Potvin O, Duchesne S. Bias-adjustment in neuroimaging-based brain age frameworks: A robust scheme. NeuroImage: Clinical 24, (2019).

30. Gaser C, Kalc P, Cole JH. A perspective on brain-age estimation and its clinical promise. Nat Comput Sci 4, 744–751 (2024).

31. Butler ER, et al. Pitfalls in brain age analyses. Hum Brain Mapp 42, 4092–4101 (2021).

32. Xiao X, et al. Brain Functional Connectome Defines a Transdiagnostic Dimension Shared by Cognitive Function and Psychopathology in Preadolescents. Biological Psychiatry, (2023).

33. Yarkoni T, Poldrack RA, Nichols TE, Van Essen DC, Wager TD. Large-scale automated synthesis of human functional neuroimaging data. Nat Methods 8, 665–670 (2011).

34. Yeo BT, et al. The organization of the human cerebral cortex estimated by intrinsic functional connectivity. J Neurophysiol 106, 1125–1165 (2011).

35. Yeh FC. Population-based tract-to-region connectome of the human brain and its hierarchical topology. Nat Commun 13, 4933 (2022).

36. Carozza S, Kletenik I, Astle D, Schwamm L, Dhand A. Whole-brain white matter variation across childhood environments. Proc Natl Acad Sci U S A 122, e2409985122 (2025).

37. Yeatman JD, Wandell BA, Mezer AA. Lifespan maturation and degeneration of human brain white matter. Nat Commun 5, 4932 (2014).

38. Bagautdinova J, et al. Development of white matter fiber covariance networks supports executive function in youth. Cell Reports 42, (2023).

39. Richie-Halford A, Yeatman J, Simon N, Rokem A. Multidimensional analysis and detection of informative features in human brain white matter. Plos Computational Biology 17, (2021).

40. Drobinin V, Van Gestel H, Helmick CA, Schmidt MH, Bowen CV, Uher R. The Developmental Brain Age Is Associated With Adversity, Depression, and Functional Outcomes Among Adolescents. Biological Psychiatry: Cognitive Neuroscience and Neuroimaging 7, 406–414 (2022).

41. Roy E, et al. Differences in educational opportunity predict white matter development. Dev Cogn Neurosci 67, 101386 (2024).

42. Goddings AL, Roalf D, Lebel C, Tamnes CK. Development of white matter microstructure and executive functions during childhood and adolescence: a review of diffusion MRI studies. Dev Cogn Neurosci 51, 101008 (2021).

43. Cardenas-Iniguez C, et al. Direct and Indirect Associations of Widespread Individual Differences in Brain White Matter Microstructure With Executive Functioning and General and Specific Dimensions of Psychopathology in Children. Biological Psychiatry: Cognitive Neuroscience and Neuroimaging 7, 362–375 (2022).

44. Cetin-Karayumak S, et al. White matter abnormalities across the lifespan of schizophrenia: a harmonized multi-site diffusion MRI study. Mol Psychiatry 25, 3208–3219 (2020).

45. Kochunov P, Hong LE. Neurodevelopmental and Neurodegenerative Models of Schizophrenia: White Matter at the Center Stage. Schizophrenia Bulletin 40, 721–728 (2014).

46. Keller AS, et al. Personalized functional brain network topography is associated with individual differences in youth cognition. Nature Communications 14, (2023).

47. Segal A, et al. Regional, circuit and network heterogeneity of brain abnormalities in psychiatric disorders. Nat Neurosci 26, 1613–1629 (2023).

48. Johnson MB, et al. Functional and evolutionary insights into human brain development through global transcriptome analysis. Neuron 62, 494–509 (2009).

49. Menon V. Large-scale brain networks and psychopathology: a unifying triple network model. Trends Cogn Sci 15, 483–506 (2011).

50. Luna B, Marek S, Larsen B, Tervo-Clemmens B, Chahal R. An integrative model of the maturation of cognitive control. Annu Rev Neurosci 38, 151–170 (2015).

51. Chaddock-Heyman L, et al. Scholastic performance and functional connectivity of brain networks in children. PLoS One 13, e0190073 (2018).

52. Wang D, Fan Q, Xiao X, He H, Yang Y, Li Y. Structural Fingerprinting of the Frontal Aslant Tract: Predicting Cognitive Control Capacity and Obsessive-Compulsive Symptoms. J Neurosci 43, 7016–7027 (2023).

53. Simmonds DJ, Hallquist MN, Asato M, Luna B. Developmental stages and sex differences of white matter and behavioral development through adolescence: A longitudinal diffusion tensor imaging (DTI) study. Neuroimage 92, 356–368 (2014).

54. Taylor JJ, et al. A transdiagnostic network for psychiatric illness derived from atrophy and lesions. Nat Hum Behav 7, 420–429 (2023).

55. Kochunov P, et al. Fractional anisotropy of water diffusion in cerebral white matter across the lifespan. Neurobiol Aging 33, 9–20 (2012).

56. Meyer HC, Lee FS. Translating Developmental Neuroscience to Understand Risk for Psychiatric Disorders. American Journal of Psychiatry 176, 179–185 (2019).

57. Mattson MP, Arumugam TV. Hallmarks of Brain Aging: Adaptive and Pathological Modification by Metabolic States. Cell Metab 27, 1176–1199 (2018).

58. Casimir P, Iwata R, Vanderhaeghen P. Linking mitochondria metabolism, developmental timing, and human brain evolution. Curr Opin Genet Dev 86, 102182 (2024).

59. van den Ameele J, et al. [(11)C]PK11195-PET Brain Imaging of the Mitochondrial Translocator Protein in Mitochondrial Disease. Neurology 96, e2761–e2773 (2021).

60. Kumar A, Muzik O, Shandal V, Chugani D, Chakraborty P, Chugani HT. Evaluation of age-related changes in translocator protein (TSPO) in human brain using (11)C-[R]- PK11195 PET. J Neuroinflammation 9, 232 (2012).

61. Somerville LH, et al. The Lifespan Human Connectome Project in Development: A large-scale study of brain connectivity development in 5-21 year olds. Neuroimage 183, 456–468 (2018).

62. Alexander LM, et al. An open resource for transdiagnostic research in pediatric mental health and learning disorders. Sci Data 4, 170181 (2017).

63. Casey BJ, et al. The Adolescent Brain Cognitive Development (ABCD) study: Imaging acquisition across 21 sites. Developmental Cognitive Neuroscience 32, 43–54 (2018).

64. Glasser MF, et al. The minimal preprocessing pipelines for the Human Connectome Project. Neuroimage 80, 105–124 (2013).

65. Richie-Halford A, et al. An analysis-ready and quality controlled resource for pediatric brain white-matter research. Sci Data 9, 616 (2022).

66. Meisler SL, Gabrieli JDE. Fiber-specific structural properties relate to reading skills in children and adolescents. Elife 11, (2022).

67. Hagler DJ, et al. Image processing and analysis methods for the Adolescent Brain Cognitive Development Study. NeuroImage 202, (2019).

68. Achenbach TM, Verhulst F. Achenbach system of empirically based assessment (ASEBA). *Burlington*, Vermont, (2010).

69. Zapolski TC, Stairs AM, Settles RF, Combs JL, Smith GT. The measurement of dispositions to rash action in children. Assessment 17, 116–125 (2010).

70. Barch DM, et al. Demographic, physical and mental health assessments in the adolescent brain and cognitive development study: Rationale and description. Dev Cogn Neurosci 32, 55–66 (2018).

71. Loewy RL, Pearson R, Vinogradov S, Bearden CE, Cannon TD. Psychosis risk screening with the Prodromal Questionnaire — Brief Version (PQ-B). Schizophrenia Research 129, 42–46 (2011).

72. Karcher NR, et al. Assessment of the Prodromal Questionnaire–Brief Child Version for Measurement of Self-reported Psychoticlike Experiences in Childhood. JAMA Psychiatry 75, (2018).

73. Fang-Cheng Y, Wedeen VJ, Tseng W-YI. Generalized ${ q}$-Sampling Imaging. IEEE Transactions on Medical Imaging 29, 1626–1635 (2010).

74. Yeh FC, Verstynen TD, Wang YB, Fernández-Miranda JC, Tseng WYI. Deterministic Diffusion Fiber Tracking Improved by Quantitative Anisotropy. Plos One 8, (2013).

75. Yeh F-C. Shape analysis of the human association pathways. NeuroImage 223, (2020).

76. Yeh F-C, et al. Automatic Removal of False Connections in Diffusion MRI Tractography Using Topology-Informed Pruning (TIP). Neurotherapeutics 16, 52–58 (2019).

77. Fortin JP, et al. Harmonization of multi-site diffusion tensor imaging data. Neuroimage 161, 149–170 (2017).

78. Xia CH, et al. Linked dimensions of psychopathology and connectivity in functional brain networks. Nat Commun 9, 3003 (2018).

79. Smolker HR, et al. The Emotional Word-Emotional Face Stroop task in the ABCD study: Psychometric validation and associations with measures of cognition and psychopathology. Dev Cogn Neurosci 53, 101054 (2022).

80. Petersen AC, Crockett L, Richards M, Boxer A. A self-report measure of pubertal status: Reliability, validity, and initial norms. Journal of Youth and Adolescence 17, 117–133 (1988).

81. Rasmussen AR, et al. Validity of self-assessment of pubertal maturation. Pediatrics 135, 86–93 (2015).

82. Kraft D, Alnaes D, Kaufmann T. Domain adapted brain network fusion captures variance related to pubertal brain development and mental health. Nat Commun 14, 6698 (2023).

83. Burt JB, Helmer M, Shinn M, Anticevic A, Murray JD. Generative modeling of brain maps with spatial autocorrelation. Neuroimage 220, 117038 (2020).

84. Vasa F, et al. Adolescent Tuning of Association Cortex in Human Structural Brain Networks. Cereb Cortex 28, 281–294 (2018).

