## Supplemental Material for "Maturation of Dorsal Association Tracts during Preadolescence Links to Concurrent and Future Cognitive Performance and Transdiagnostic Psychopathology"

### Supplementary Material

**Table S1. Names, abbreviations and mainly connected brain regions (DK atlas) of the included white matter (WM) tracts**

| Tract Name | Abbrev. | Region 1 | Region 2 |
|---|---|---|---|
| Arcuate Fasciculus | AF | Superior temporal | Pars opercularis |
| Frontal Aslant Tract | FAT | Superior frontal | Pars opercularis |
| Parietal Aslant Tract | PAT | Inferior parietal | Middle temporal |
| Superior Longitudinal Fasciculus 1 | SLF1 | Superior frontal | Precuneus |
| Superior Longitudinal Fasciculus 2 | SLF2 | Inferior parietal | Precentral |
| Superior Longitudinal Fasciculus 3 | SLF3 | Inferior parietal | Pars opercularis |
| Inferior Fronto Occipital Fasciculus | IFOF | Lateral orbitofrontal | Lateral occipital |
| Inferior Longitudinal Fasciculus | ILF | Lateral occipital | Superior temporal |
| Uncinate Fasciculus | UF | Superior temporal | Lateral orbitofrontal |
| Middle Longitudinal Fasciculus | MdLF | Superior temporal | Middle temporal |
| Cingulum Frontal Parahippocampal | C_FPH | Isthmus cingulate | Superior frontal |
| Cingulum Frontal Parietal | C_FP | Precuneus | Rostral anterior cingulate |
| Cingulum Parahippocampal | C_PH | Isthmus cingulate | Inferior temporal |
| Cingulum Parahippocampal Parietal | C_PHP | Entorhinal | Precuneus |
| Cingulum Parolfactory | C_PO | Medial orbitofrontal | Precuneus |
| Fornix | F | Mammillary body | Hippocampus |
| Corticospinal Tract | CST | Precentral | Brainstem |
| Optic Radiation | OR | Lateral occipital | Thalamus |
| Vertical Occipital Fasciculus | VOF | Superior parietal | Lateral occipital |
| Thalamic Radiation Anterior | TR_A | Thalamus | Lateral orbitofrontal |
| Thalamic Radiation Posterior | TR_P | Thalamus | Lateral occipital |
| Thalamic Radiation Superior | TR_S | Thalamus | Superior parietal |
| Corticostriatal Tract Anterior | CS_A | Striatum | Lateral orbitofrontal/ Superior frontal |
| Corticostriatal Tract Posterior | CS_P | Striatum | Pericalcarine/Inferior_parietal/ Lateral occipital |
| Corticostriatal Tract Superior | CS_S | Striatum | Superior parietal/ Precentral/Postcentral/Superiorfrontal /Paracentral/Postcentral |
| Corpus Callosum Body | CC_Body | Superior frontal/Paracentral/Postcentral/Superior parietal | |
| Corpus Callosum Forceps Major | CC_Major | Cuneus/Superior parietal | |
| Corpus Callosum Forceps Minor | CC_Minor | Frontalpole/Superior frontal/Medial orbitofrontal | |
| Corpus Callosum Tapetum | CC_Tap | Hippocampus | |

Notes: the connected cortical regions are defined in the Desikan-Killiany (DK) atlas[1].

**Table S2. Five-fold cross-validation performance of prediction models in HCP-D dataset**

| Tract | Uncorrected | | Corrected | |
|---|---|---|---|---|
| | $R^2$ | MAE (years) | $R^2$ | MAE (years) |
| AF | 0.397 | 2.46 | 0.797 | 1.54 |
| C_FPH | 0.431 | 2.49 | 0.809 | 1.52 |
| C_FP | 0.393 | 2.57 | 0.808 | 1.54 |
| C_PH | 0.371 | 2.59 | 0.804 | 1.51 |
| C_PHP | 0.396 | 2.54 | 0.802 | 1.52 |
| C_PO | 0.268 | 2.85 | 0.836 | 1.38 |
| CST | 0.502 | 2.27 | 0.801 | 1.50 |
| CS_A | 0.357 | 2.61 | 0.814 | 1.44 |
| CS_P | 0.556 | 2.12 | 0.804 | 1.46 |
| CS_S | 0.511 | 2.30 | 0.799 | 1.53 |
| F | 0.240 | 2.87 | 0.842 | 1.38 |
| FAT | 0.538 | 2.18 | 0.800 | 1.51 |
| IFOF | 0.436 | 2.42 | 0.805 | 1.47 |
| ILF | 0.397 | 2.56 | 0.800 | 1.47 |
| MdLF | 0.443 | 2.44 | 0.807 | 1.49 |
| OR | 0.393 | 2.53 | 0.806 | 1.51 |
| PAT | 0.576 | 2.09 | 0.811 | 1.52 |
| SLF1 | 0.416 | 2.45 | 0.803 | 1.54 |
| SLF2 | 0.497 | 2.27 | 0.799 | 1.53 |
| SLF3 | 0.470 | 2.34 | 0.810 | 1.46 |
| TR_A | 0.521 | 2.26 | 0.813 | 1.52 |
| TR_P | 0.547 | 2.17 | 0.805 | 1.53 |
| TR_S | 0.518 | 2.26 | 0.812 | 1.51 |
| UF | 0.242 | 2.87 | 0.840 | 1.35 |
| VOF | 0.426 | 2.45 | 0.804 | 1.54 |
| CC_Body | 0.609 | 2.46 | 0.828 | 1.54 |
| CC_Major | 0.463 | 2.49 | 0.824 | 1.52 |
| CC_Minor | 0.169 | 2.57 | 0.875 | 1.54 |
| CC_Tap | 0.363 | 2.59 | 0.824 | 1.51 |

**Table S3. Testing performance of brain age prediction models in HBN dataset**

| Tract | Uncorrected | | Corrected | |
|---|---|---|---|---|
| | $R^2$ | MAE (years) | $R^2$ | MAE (years) |
| Whole-brain | 0.387 | 2.798 | 0.694 | 1.651 |
| AF | 0.053 | 3.752 | 0.528 | 2.072 |
| C_FPH | 0.140 | 3.430 | 0.558 | 2.103 |
| C_FP | 0.148 | 3.391 | 0.676 | 1.648 |
| C_PH | 0.102 | 3.462 | 0.697 | 1.618 |
| C_PHP | 0.200 | 3.277 | 0.703 | 1.570 |
| C_PO | 0.042 | 3.879 | 0.736 | 1.462 |
| CST | 0.112 | 3.550 | 0.524 | 2.163 |
| CS_A | 0.091 | 4.285 | 0.314 | 3.510 |
| CS_P | 0.218 | 3.172 | 0.618 | 1.898 |
| CS_S | 0.300 | 2.842 | 0.609 | 1.896 |
| F | 0.084 | 3.717 | 0.775 | 1.352 |
| FAT | 0.299 | 2.929 | 0.632 | 1.909 |
| IFOF | 0.137 | 3.489 | 0.585 | 2.021 |
| ILF | 0.107 | 3.578 | 0.632 | 1.830 |
| MdLF | 0.098 | 3.620 | 0.565 | 2.050 |
| OR | 0.294 | 3.143 | 0.746 | 1.498 |
| PAT | 0.006 | 4.130 | 0.288 | 2.744 |
| SLF1 | 0.128 | 3.475 | 0.625 | 1.802 |
| SLF2 | 0.144 | 3.519 | 0.483 | 2.358 |
| SLF3 | 0.023 | 4.067 | 0.410 | 2.414 |
| TR_A | 0.213 | 3.083 | 0.585 | 1.981 |
| TR_P | 0.312 | 3.011 | 0.693 | 1.641 |
| TR_S | 0.191 | 3.278 | 0.491 | 2.381 |
| UF | 0.015 | 3.974 | 0.631 | 1.741 |
| VOF | 0.067 | 3.683 | 0.615 | 1.842 |
| CC_Body | 0.342 | 2.913 | 0.646 | 1.963 |
| CC_Major | 0.208 | 3.372 | 0.683 | 1.742 |
| CC_Minor | 0.023 | 4.069 | 0.686 | 1.644 |
| CC_Tap | 0.290 | 3.149 | 0.779 | 1.354 |

**Table S4. Testing performance of brain age prediction models in independent ABCD dataset**

| Tract | Uncorrected | | Corrected | |
|---|---|---|---|---|
| | $R^2$ | MAE (years) | $R^2$ | MAE (years) |
| AF | 0.051 | 2.69 | 0.187 | 2.03 |
| C_FPH | 0.065 | 2.37 | 0.191 | 1.84 |
| C_FP | 0.086 | 2.22 | 0.255 | 1.57 |
| C_PH | 0.064 | 2.40 | 0.266 | 1.50 |
| C_PHP | 0.059 | 2.39 | 0.245 | 1.50 |
| C_PO | 0.033 | 2.77 | 0.261 | 1.50 |
| CST | 0.094 | 2.17 | 0.262 | 1.66 |
| CS_A | 0.022 | 3.74 | 0.076 | 3.33 |
| CS_P | 0.112 | 2.03 | 0.248 | 1.64 |
| CS_S | 0.115 | 2.04 | 0.239 | 1.61 |
| F | 0.026 | 2.71 | 0.263 | 1.40 |
| FAT | 0.060 | 2.62 | 0.176 | 2.16 |
| IFOF | 0.075 | 2.28 | 0.242 | 1.72 |
| ILF | 0.062 | 2.50 | 0.196 | 2.04 |
| MdLF | 0.058 | 2.53 | 0.230 | 1.81 |
| OR | 0.055 | 2.49 | 0.250 | 1.51 |
| PAT | 0.023 | 3.50 | 0.125 | 2.69 |
| SLF1 | 0.104 | 2.29 | 0.246 | 1.76 |
| SLF2 | 0.059 | 3.19 | 0.117 | 2.97 |
| SLF3 | 0.044 | 3.19 | 0.120 | 2.80 |
| TR_A | 0.116 | 2.07 | 0.205 | 1.81 |
| TR_P | 0.078 | 2.30 | 0.217 | 1.81 |
| TR_S | 0.057 | 2.64 | 0.113 | 2.46 |
| UF | 0.032 | 2.89 | 0.158 | 2.10 |
| VOF | 0.009 | 3.33 | 0.081 | 2.74 |
| CC_Body | 0.079 | 2.42 | 0.169 | 2.19 |
| CC_Major | 0.058 | 2.62 | 0.126 | 2.45 |
| CC_Minor | 0.014 | 3.20 | 0.248 | 1.63 |
| CC_Tap | 0.033 | 2.60 | 0.261 | 1.36 |

**Table S5. Comparisons of loading distributions from two sCCA modes derived from bootstrap resampling tests (N = 1,000)**

| Behavioral measure | T | Effect size (Cohen's *d*) |
|---|---|---|
| CBCL_syn_social_prob | 3.770 | 0.158 |
| **CBCL_syn_attention_prob** | **-16.773** | **0.927** |
| **CBCL_total** | **-25.241** | **1.387** |
| **CBCL_dsm_ADHD** | **-23.537** | **1.269** |
| **PGBI_mania_total** | **24.463** | **0.920** |
| nihtbx_pic_vocab | -3.117 | 0.205 |
| nihtbx_flanker | -7.195 | 0.332 |
| nihtbx_list_sort | 6.543 | 0.379 |
| **nihtbx_card_sort** | **-10.869** | **0.668** |
| **nihtbx_pattern_comp** | -17.167 | **1.010** |
| nihtbx_pic_seq_memory | -6.261 | 0.264 |
| nihtbx_oral_reading | 3.089 | 0.185 |
| nihtbx_fluid_comp | -0.725 | 0.053 |
| nihtbx_cryst_comp | 1.859 | 0.130 |
| nihtbx_total_comp | 1.627 | 0.124 |
| **little_man_efficiency** | **-20.694** | **1.074** |
| matrix_reasoning | 1.558 | 0.086 |
| verbal_learning_immediate | -6.073 | 0.234 |
| verbal_learning_delayed | -0.991 | 0.035 |
| **SST_ssrt_reversed** | **28.713** | **0.977** |
| X2_back | -1.569 | 0.091 |
| recog_memory | -1.024 | 0.049 |

**Notes**: Behavioral measures showing significant differences in loading distributions between the two sCCA modes (effect size > 0.5) are highlighted.

**Table S6. Selected Neurosynth topics and their abbreviations**

| Abbreviation | Topic |
|---|---|
| Attention | 49_attention_attentional_task |
| Cognitive-control and task performance | 11_task_cognitive_performance |
| Cognitive task demands | 36_task_difficulty_demands |
| Numerical Representation and Processing | 18_number_numerical_numbers |
| Representation | 55_representations_representation_abstract |
| Learning and training | 48_learning_training_sequence |
| Language | 12_language_comprehension_sentences |
| Reading and words | 21_reading_words_word |
| Working memory | 45_memory_working_wm |
| Mental imagery | 9_imagery_mental_events |
| Cognitive Conflict | 14_conflict_interference_control |
| Response inhibition | 89_inhibition_response_control |
| Contextual Modulation and Influence | 42_context_influence_modulation |
| Speech | 31_speech_auditory_production |
| Perception | 50_perceptual_perception_interaction |
| Semantic words | 23_semantic_words_word |
| Schizophrenia and Psychotic Symptoms | 10_schizophrenia_symptoms_risk |
| Social Cognition and Empathy | 92_social_empathy_moral |
| Recognition Memory | 76_memory_recognition_items |
| Memory Encoding | 72_memory_encoding_hippocampus |
| Reasoning | 71_reasoning_mind_mental |
| Emotion Regulation | 35_regulation_emotion_reappraisal |
| Personality Traits | 58_personality_trait_traits |
| Decision making | 53_decision_making_risk |
| Episodic Memory | 41_memory_retrieval_episodic |
| Emotion and Affective Evaluation | 30_emotional_negative_positive |
| Depression | 5_depression_mdd_disorder |
| ADHD and Eating Disorders | 54_adhd_food_weight |
| Substance Abuse and Stress Response | 75_stress_alcohol_users |
| Fear Conditioning and PTSD | 26_fear_anxiety_ptsd |
| Reward | 33_reward_striatum_ventral |

**Table S7. Functional decoding results for selected Neurosynth topics. Significant results are bolded for each tract.**

| Topic | Mode 1 | | Mode 2 | |
|---|---|---|---|---|
| | rho | $p_{FDR}$ | rho | $p_{FDR}$ |
| Depression | -0.547 | 1.000 | 0.543 | **0.037** |
| Mental imagery | 0.432 | 0.093 | -0.240 | 1.000 |
| Schizophrenia and Psychotic Symptoms | -0.138 | 1.000 | -0.033 | 1.000 |
| Cognitive-control and task performance | 0.536 | **0.037** | -0.475 | 1.000 |
| Language | 0.549 | **0.037** | -0.666 | 1.000 |
| Cognitive Conflict | 0.387 | 0.158 | -0.351 | 1.000 |
| Numerical Representation and Processing | 0.522 | **0.043** | -0.397 | 1.000 |
| Reading and Words | 0.600 | **0.037** | -0.673 | 1.000 |
| Semantic words | 0.466 | 0.053 | -0.495 | 1.000 |
| Fear Conditioning and PTSD | -0.532 | 1.000 | 0.530 | **0.037** |
| Emotion and Affective Evaluation | -0.373 | 1.000 | 0.338 | 0.217 |
| Speech | 0.612 | **0.037** | -0.706 | 1.000 |
| Reward | -0.641 | 1.000 | 0.508 | **0.041** |
| Emotion Regulation | -0.171 | 1.000 | 0.034 | 0.887 |
| Cognitive task demands | 0.596 | **0.037** | -0.647 | 1.000 |
| Episodic Memory | -0.246 | 1.000 | 0.297 | 0.230 |
| Working_memory | 0.363 | 0.217 | -0.273 | 1.000 |
| Learning and training | 0.234 | 0.437 | -0.122 | 1.000 |
| Attention | 0.606 | **0.037** | -0.540 | 1.000 |
| Perception | 0.410 | 0.128 | -0.392 | 1.000 |
| Decision making | -0.299 | 1.000 | 0.210 | 0.444 |
| ADHD and Eating Disorders | -0.533 | 1.000 | 0.514 | **0.039** |
| Representation | 0.599 | **0.037** | -0.505 | 1.000 |
| Personality Traits | -0.328 | 1.000 | 0.240 | 0.364 |
| Reasoning | 0.085 | 0.815 | -0.089 | 1.000 |
| Memory Encoding | -0.198 | 1.000 | 0.273 | 0.299 |
| OCD and Dopamine Treatment | -0.555 | 1.000 | 0.452 | 0.070 |
| Substance Abuse and Stress Response | -0.610 | 1.000 | 0.570 | **0.037** |
| Recognition Memory | -0.022 | 1.000 | 0.153 | 0.599 |
| Response inhibition | -0.017 | 1.000 | -0.012 | 1.000 |
| Social Cognition and Empathy | 0.057 | 0.887 | -0.139 | 1.000 |

**Table S8. Statistics for the relationship between baseline tract-based BAGs and school grade at 2-y-follow-up**

| Tract-based BAG | F | $\beta$ [95% CI] | $\Delta R^2_{adjusted}$ | $p_{FDR}$ |
|---|---|---|---|---|
| Whole-brain | 23.760 | -0.070 [-0.099, -0.042] | 0.286 | < 0.001 |
| AF | 41.194 | -0.054 [-0.071, -0.038] | 0.503 | < 0.001 |
| FAT | 31.234 | -0.044 [-0.060, -0.029] | 0.379 | < 0.001 |
| PAT | 52.488 | -0.040 [-0.051, -0.029] | 0.644 | < 0.001 |
| SLF2 | 44.426 | -0.036 [-0.047, -0.025] | 0.544 | < 0.001 |
| SLF3 | 33.153 | -0.033 [-0.045, -0.022] | 0.403 | < 0.001 |
| SLF1 | 37.310 | -0.059 [-0.078, -0.040] | 0.455 | < 0.001 |
| IFOF | 7.415 | -0.030 [-0.051, -0.008] | 0.081 | 0.008 |
| ILF | 16.250 | -0.035 [-0.051, - 0.018] | 0.192 | < 0.001 |
| UF | 5.560 | -0.019 [-0.035, -0.003] | 0.057 | 0.022 |
| MdLF | 41.015 | -0.062 [-0.080, -0.043] | 0.501 | < 0.001 |
| VOF | 8.536 | -0.019 [-0.031, -0.006] | 0.095 | 0.005 |
| OR | 11.883 | -0.043 [-0.067, -0.019] | 0.137 | 0.001 |
| CST | 17.278 | -0.046 [-0.067, -0.024] | 0.205 | < 0.001 |
| C_PHP | 10.268 | -0.039 [-0.063, -0.015] | 0.117 | 0.002 |
| C_PH | 5.628 | -0.029 [-0.053, -0.005] | 0.058 | 0.022 |
| C_FP | 26.583 | -0.061 [-0.084, -0.038] | 0.321 | < 0.001 |
| C_FPH | 0.001 | 0.000 [-0.021, 0.020] | -0.013 | 0.974 |
| C_PO | 1.614 | -0.016 [-0.039, 0.008] | 0.008 | 0.218 |
| F | 25.278 | -0.065 [-0.090, -0.040] | 0.305 | < 0.001 |
| CS_A | 0.724 | -0.004 [-0.014, 0.006] | -0.003 | 0.408 |
| CS_P | 13.466 | -0.039 [-0.060, -0.018] | 0.157 | < 0.001 |
| CS_S | 8.158 | -0.032 [-0.053, -0.010] | 0.090 | 0.006 |
| TR_A | 20.614 | -0.046 [-0.066, -0.026] | 0.246 | < 0.001 |
| TR_P | 22.980 | -0.046 [-0.065, -0.027] | 0.276 | < 0.001 |
| TR_S | 2.758 | -0.011 [-0.024, 0.002] | 0.022 | 0.107 |
| CC_Body | 8.981 | -0.024 [-0.040, -0.008] | 0.100 | 0.004 |
| CC_Major | 8.817 | -0.021 [-0.035, -0.007] | 0.098 | 0.004 |
| CC_Minor | 17.030 | -0.047 [-0.069, -0.025] | 0.201 | < 0.001 |
| CC_Tap | 4.154 | -0.027 [-0.053, -0.001] | 0.040 | 0.048 |

**Table S9. Statistics for the relationship between baseline tract-based BAGs and math ability (assessed via Stanford Mental Arithmetic Response Time Evaluation [SAMRTE]) at 3-y-follow-up**

| Tract-based BAG | F | $\beta$ [95% CI] | $\Delta R^2_{adjusted}$ | $p_{FDR}$ |
|---|---|---|---|---|
| Whole-brain | 64.320 | 1.519 [1.148, 1.890] | 1.063 | < 0.001 |
| AF | 49.573 | 0.792 [0.572, 1.013] | 0.817 | < 0.001 |
| FAT | 68.074 | 0.863 [0.658, 1.068] | 1.125 | < 0.001 |
| PAT | 53.814 | 0.542 [0.397, 0.687] | 0.888 | < 0.001 |
| SLF2 | 47.380 | 0.495 [0.354, 0.636] | 0.781 | < 0.001 |
| SLF3 | 44.205 | 0.512 [0.361, 0.663] | 0.728 | < 0.001 |
| SLF1 | 35.552 | 0.759 [0.509, 1.008] | 0.583 | < 0.001 |
| IFOF | 46.138 | 0.966 [0.687, 1.244] | 0.760 | < 0.001 |
| ILF | 54.206 | 0.828 [0.608, 1.049] | 0.895 | < 0.001 |
| UF | 19.340 | 0.471 [0.261, 0.682] | 0.310 | < 0.001 |
| MdLF | 36.188 | 0.766 [0.516, 1.015] | 0.594 | < 0.001 |
| VOF | 5.809 | 0.203 [0.038, 0.368] | 0.082 | 0.016 |
| OR | 35.647 | 0.981 [0.659, 1.303] | 0.585 | < 0.001 |
| CST | 32.009 | 0.819 [0.536, 1.103] | 0.523 | < 0.001 |
| C_PHP | 34.129 | 0.938 [0.623, 1.253] | 0.559 | < 0.001 |
| C_PH | 16.803 | 0.669 [0.349, 0.989] | 0.267 | < 0.001 |
| C_FP | 23.020 | 0.757 [0.448, 1.066] | 0.372 | < 0.001 |
| C_FPH | 1.832 | 0.184 [-0.083, 0.451] | 0.014 | 0.176 |
| C_PO | 6.847 | 0.423 [0.106, 0.740] | 0.099 | 0.010 |
| F | 32.605 | 0.971 [0.638, 1.305] | 0.533 | < 0.001 |
| CS_A | 15.221 | 0.262 [0.131, 0.394] | 0.241 | < 0.001 |
| CS_P | 89.388 | 1.310 [1.039, 1.582] | 1.477 | < 0.001 |
| CS_S | 32.744 | 0.833 [0.547, 1.118] | 0.536 | < 0.001 |
| TR_A | 48.453 | 0.922 [0.663, 1.182] | 0.799 | < 0.001 |
| TR_P | 44.023 | 0.843 [0.594, 1.092] | 0.725 | < 0.001 |
| TR_S | 6.364 | 0.214 [0.048, 0.380] | 0.091 | < 0.001 |
| CC_Body | 28.155 | 0.566 [0.357, 0.775] | 0.459 | < 0.001 |
| CC_Major | 28.701 | 0.506 [0.321, 0.691] | 0.468 | < 0.001 |
| CC_Minor | 19.521 | 0.669 [0.372, 0.966] | 0.313 | < 0.001 |
| CC_Tap | 27.416 | 0.927 [0.580, 1.274] | 0.446 | < 0.001 |

**Table S10. Statistics for the relationship between baseline tract-based BAGs and overall accuracy of emotional Stroop task at 3-y-follow-up**

| Tract-based BAG | F | $\beta$ [95% CI] | $\Delta R^2_{adjusted}$ | $p_{FDR}$ |
|---|---|---|---|---|
| Whole-brain | 26.364 | 0.067 [0.041, 0.092] | 0.418 | < 0.001 |
| AF | 16.615 | 0.019 [0.010, 0.029] | 0.258 | < 0.001 |
| FAT | 38.334 | 0.027 [0.019, 0.036] | 0.614 | < 0.001 |
| PAT | 22.028 | 0.015 [0.009, 0.021] | 0.347 | < 0.001 |
| SLF2 | 23.496 | 0.015 [0.009, 0.021] | 0.371 | < 0.001 |
| SLF3 | 31.577 | 0.018 [0.012, 0.025] | 0.504 | < 0.001 |
| SLF1 | 11.657 | 0.018 [0.008, 0.029] | 0.176 | 0.002 |
| IFOF | 5.801 | 0.015 [0.003, 0.026] | 0.079 | 0.026 |
| ILF | 14.952 | 0.018 [0.009, 0.028] | 0.230 | < 0.001 |
| UF | 4.973 | 0.010 [0.001, 0.019] | 0.066 | 0.038 |
| MdLF | 15.459 | 0.021 [0.011, 0.032] | 0.239 | < 0.001 |
| VOF | 1.895 | 0.005 [-0.002, 0.012] | 0.015 | 0.193 |
| OR | 0.208 | 0.003 [-0.010, 0.017] | -0.013 | 0.670 |
| CST | 11.263 | 0.021 [0.009, 0.032] | 0.170 | 0.002 |
| C_PHP | 5.597 | 0.016 [0.003, 0.029] | 0.076 | 0.027 |
| C_PH | 7.269 | 0.019 [0.005, 0.032] | 0.104 | 0.014 |
| C_FP | 3.403 | 0.012 [-0.001, 0.025] | 0.040 | 0.085 |
| C_FPH | 0.024 | 0.001 [-0.010, 0.012] | -0.016 | 0.876 |
| C_PO | 6.805 | 0.018 [0.004, 0.031] | 0.096 | 0.017 |
| F | 6.307 | 0.018 [0.004, 0.032] | 0.088 | 0.021 |
| CS_A | 1.092 | 0.003 [-0.003, 0.009] | 0.002 | 0.327 |
| CS_P | 19.941 | 0.026 [0.015, 0.038] | 0.313 | < 0.001 |
| CS_S | 3.188 | 0.011 [-0.001, 0.023] | 0.036 | 0.091 |
| TR_A | 8.380 | 0.016 [0.005, 0.027] | 0.122 | 0.008 |
| TR_P | 15.838 | 0.021 [0.011, 0.032] | 0.245 | < 0.001 |
| TR_S | 0.650 | -0.003 [-0.010, 0.004] | -0.006 | 0.448 |
| CC_Body | 5.771 | 0.011 [0.002, 0.020] | 0.079 | 0.026 |
| CC_Major | 4.718 | 0.009 [0.001, 0.017] | 0.062 | 0.042 |
| CC_Minor | 3.375 | 0.012 [-0.001, 0.024] | 0.039 | 0.085 |
| CC_Tap | 2.290 | 0.011 [-0.003, 0.026] | 0.021 | 0.154 |

**Table S11**. GLM analysis results on the relationships between 2-y-follow-up tract-BAG measures (with age-bias correction) and 2-y-follow-up transdiagnostic status.

| Tract-based measure | F | $p_{FDR}$ |
|---|---|---|
| Association Development Mode | 7.13 | 0.010 |
| Subcortical/Limbic Development Mode | 4.95 | 0.033 |
| Whole-brain | 3.85 | 0.066 |
| AF | 2.72 | 0.140 |
| FAT | 5.99 | 0.020 |
| PAT | 7.01 | 0.010 |
| SLF2 | 1.90 | 0.219 |
| SLF3 | 3.24 | 0.093 |
| SLF1 | 8.93 | 0.004 |
| IFOF | 2.37 | 0.177 |
| ILF | 1.89 | 0.219 |
| UF | 0.44 | 0.708 |
| MdLF | 4.51 | 0.039 |
| VOF | 3.21 | 0.093 |
| OR | 1.64 | 0.249 |
| CST | 5.10 | 0.033 |
| C_PHP | 2.55 | 0.156 |
| C_PH | 2.22 | 0.193 |
| C_FP | 5.41 | 0.029 |
| C_FPH | 0.88 | 0.476 |
| C_PO | 1.94 | 0.219 |
| F | 1.56 | 0.260 |
| CS_A | 0.19 | 0.825 |
| CS_P | 1.71 | 0.246 |
| CS_S | 1.69 | 0.246 |
| TR_A | 4.58 | 0.039 |
| TR_P | 3.34 | 0.093 |
| TR_S | 0.38 | 0.718 |
| CC_Body | 3.78 | 0.066 |
| CC_Major | 1.94 | 0.219 |
| CC_Minor | 0.36 | 0.718 |
| CC_Tap | 0.89 | 0.476 |

**Table S12**. GLM analysis results on the relationships between baseline tract-BAG measures (with age-bias correction) and the cumulative number of KSADS-5 at baseline, at 2-y-follow-up, and the transition of the transdiagnostic status between baseline and 2-y-follow-up.

| Tract-based measure | $BAG_{Baseline}$ ~ $KSADS_{baseline}$ | | $BAG_{Baseline}$ ~ $KSADS_{followup}$ | | $BAG_{Baseline}$ ~ $KSADS_{conversion}$ | |
|---|---|---|---|---|---|---|
| | F | $p_{FDR}$ | F | $p_{FDR}$ | F | $p_{FDR}$ |
| Whole-brain | 7.05 | 0.004 | 7.03 | 0.005 | 6.06 | 0.002 |
| AF | 9.11 | 0.001 | 4.29 | 0.038 | 7.04 | 0.001 |
| FAT | 9.78 | 0.001 | 8.64 | 0.002 | 7.27 | 0.001 |
| PAT | 13.89 | < 0.001 | 8.44 | 0.002 | 8.58 | < 0.001 |
| SLF2 | 8.21 | 0.002 | 5.24 | 0.020 | 4.30 | 0.014 |
| SLF3 | 5.13 | 0.022 | 2.15 | 0.175 | 4.43 | 0.014 |
| SLF1 | 7.49 | 0.003 | 8.75 | 0.002 | 8.04 | < 0.001 |
| IFOF | 5.52 | 0.017 | 3.88 | 0.048 | 2.71 | 0.078 |
| ILF | 3.14 | 0.100 | 1.76 | 0.247 | 0.87 | 0.484 |
| UF | 1.27 | 0.400 | 0.76 | 0.503 | 0.88 | 0.484 |
| MdLF | 3.90 | 0.061 | 2.46 | 0.142 | 4.35 | 0.014 |
| VOF | 1.73 | 0.295 | 0.84 | 0.498 | 1.83 | 0.187 |
| OR | 0.60 | 0.660 | 2.93 | 0.094 | 2.39 | 0.102 |
| CST | 3.69 | 0.064 | 5.32 | 0.020 | 3.11 | 0.054 |
| C_PHP | 0.52 | 0.689 | 2.26 | 0.165 | 1.65 | 0.226 |
| C_PH | 0.93 | 0.514 | 1.05 | 0.436 | 0.95 | 0.473 |
| C_FP | 3.67 | 0.064 | 4.74 | 0.026 | 3.53 | 0.038 |
| C_FPH | 0.33 | 0.770 | 0.64 | 0.545 | 0.32 | 0.811 |
| C_PO | 2.23 | 0.190 | 1.17 | 0.403 | 1.83 | 0.187 |
| F | 1.17 | 0.425 | 4.93 | 0.024 | 4.61 | 0.013 |
| CS_A | 2.27 | 0.190 | 0.33 | 0.718 | 2.46 | 0.097 |
| CS_P | 2.53 | 0.163 | 7.91 | 0.003 | 3.19 | 0.052 |
| CS_S | 0.37 | 0.765 | 0.76 | 0.503 | 1.27 | 0.347 |
| TR_A | 0.73 | 0.603 | 3.49 | 0.061 | 2.04 | 0.154 |
| TR_P | 1.61 | 0.315 | 6.15 | 0.011 | 2.60 | 0.085 |
| TR_S | 0.21 | 0.837 | 0.91 | 0.483 | 1.06 | 0.430 |
| CC_Body | 4.04 | 0.059 | 3.25 | 0.073 | 2.96 | 0.061 |
| CC_Major | 1.30 | 0.400 | 4.08 | 0.043 | 2.92 | 0.061 |
| CC_Minor | 2.51 | 0.163 | 3.57 | 0.060 | 3.48 | 0.038 |
| CC_Tap | 0.00 | 0.997 | 1.58 | 0.281 | 0.39 | 0.786 |

**Table S13**. Sensitivity analysis on the impact of age-bias correction on the relationships between baseline tract-BAG measures (without age-bias correction) and the cumulative number of KSADS-5 diagnoses at baseline, at 2-y-follow-up, and the transition of the transdiagnostic status between baseline and 2-y-follow-up.

| Tract-based measure | $BAG_{Baseline}$ ~ $KSADS_{baseline}$ | | $BAG_{Baseline}$ ~ $KSADS_{followup}$ | | $BAG_{Baseline}$ ~ $KSADS_{conversion}$ | |
|---|---|---|---|---|---|---|
| | F | $p_{FDR}$ | F | $p_{FDR}$ | F | $p_{FDR}$ |
| Association Development Mode | 15.06 | < 0.001 | 11.90 | < 0.001 | 10.53 | < 0.001 |
| Subcortical/Limbic Development Mode | 6.22 | 0.008 | 7.20 | 0.004 | 4.76 | 0.012 |
| Whole-brain | 6.36 | 0.004 | 6.70 | 0.006 | 5.42 | 0.005 |
| AF | 8.35 | 0.001 | 3.87 | 0.048 | 6.41 | 0.002 |
| FAT | 9.15 | 0.001 | 8.51 | 0.002 | 6.77 | 0.001 |
| PAT | 13.21 | < 0.001 | 8.30 | 0.002 | 8.07 | < 0.001 |
| SLF2 | 7.91 | 0.002 | 5.21 | 0.018 | 4.10 | 0.023 |
| SLF3 | 4.79 | 0.019 | 2.00 | 0.205 | 4.15 | 0.023 |
| SLF1 | 6.86 | 0.003 | 8.17 | 0.002 | 7.38 | 0.001 |
| IFOF | 4.96 | 0.014 | 3.45 | 0.068 | 2.30 | 0.127 |
| ILF | 2.78 | 0.093 | 1.73 | 0.236 | 0.69 | 0.594 |
| UF | 1.23 | 0.389 | 0.48 | 0.637 | 0.86 | 0.526 |
| MdLF | 3.32 | 0.054 | 2.19 | 0.179 | 3.77 | 0.030 |
| VOF | 1.51 | 0.283 | 0.80 | 0.525 | 1.62 | 0.253 |
| OR | 0.38 | 0.652 | 2.49 | 0.139 | 1.95 | 0.182 |
| CST | 3.21 | 0.058 | 5.36 | 0.017 | 2.80 | 0.083 |
| C_PHP | 0.29 | 0.682 | 1.80 | 0.230 | 1.23 | 0.366 |
| C_PH | 0.93 | 0.505 | 0.84 | 0.525 | 0.72 | 0.594 |
| C_FP | 3.09 | 0.058 | 4.52 | 0.032 | 2.96 | 0.083 |
| C_FPH | 0.20 | 0.767 | 0.67 | 0.563 | 0.20 | 0.893 |
| C_PO | 1.65 | 0.181 | 0.85 | 0.525 | 1.28 | 0.358 |
| F | 0.73 | 0.416 | 4.38 | 0.033 | 3.83 | 0.030 |
| CS_A | 2.07 | 0.181 | 0.27 | 0.760 | 2.30 | 0.127 |
| CS_P | 2.21 | 0.153 | 7.78 | 0.003 | 2.84 | 0.083 |
| CS_S | 0.53 | 0.759 | 0.61 | 0.577 | 1.28 | 0.358 |
| TR_A | 0.59 | 0.594 | 3.29 | 0.075 | 1.81 | 0.208 |
| TR_P | 1.33 | 0.304 | 6.13 | 0.009 | 2.24 | 0.130 |
| TR_S | 0.21 | 0.835 | 0.78 | 0.525 | 0.98 | 0.476 |
| CC_Body | 3.74 | 0.051 | 3.14 | 0.082 | 2.74 | 0.083 |
| CC_Major | 1.16 | 0.389 | 3.88 | 0.048 | 2.74 | 0.083 |
| CC_Minor | 1.89 | 0.153 | 2.65 | 0.126 | 2.68 | 0.085 |
| CC_Tap | 0.07 | 0.997 | 1.96 | 0.205 | 0.37 | 0.802 |

**Table S14**. Sensitivity analysis on the impact of including pubertal effects as additional covariate on the relationships between baseline tract-BAG measures (with age-bias correction) and the cumulative number of KSADS-5 diagnoses at baseline, at 2-y-follow-up, and the transition of the transdiagnostic status between baseline and 2-y-follow-up.

| Tract-based measure | $BAG_{Baseline}$ ~ $KSADS_{baseline}$ | | $BAG_{Baseline}$ ~ $KSADS_{followup}$ | | $BAG_{Baseline}$ ~ $KSADS_{conversion}$ | |
|---|---|---|---|---|---|---|
| | F | $p_{FDR}$ | F | $p_{FDR}$ | F | $p_{FDR}$ |
| Association Development Mode | 12.503 | < 0.001 | 10.754 | 0.001 | 9.380 | < 0.001 |
| Subcortical/Limbic Development Mode | 4.277 | 0.045 | 5.535 | 0.016 | 3.740 | 0.037 |
| Whole-brain | 5.300 | 0.023 | 5.735 | 0.016 | 5.013 | 0.008 |
| AF | 8.887 | 0.001 | 4.540 | 0.030 | 7.154 | 0.001 |
| FAT | 7.922 | 0.003 | 7.544 | 0.004 | 6.030 | 0.003 |
| PAT | 12.511 | < 0.001 | 8.110 | 0.003 | 7.918 | < 0.001 |
| SLF2 | 7.136 | 0.005 | 4.627 | 0.030 | 3.686 | 0.037 |
| SLF3 | 4.620 | 0.040 | 1.994 | 0.198 | 4.077 | 0.027 |
| SLF1 | 7.056 | 0.005 | 8.481 | 0.003 | 8.040 | < 0.001 |
| IFOF | 4.341 | 0.045 | 3.468 | 0.067 | 2.310 | 0.125 |
| ILF | 2.738 | 0.138 | 1.148 | 0.423 | 0.611 | 0.649 |
| UF | 1.396 | 0.390 | 0.478 | 0.640 | 0.771 | 0.564 |
| MdLF | 2.775 | 0.138 | 2.038 | 0.198 | 3.513 | 0.042 |
| VOF | 1.363 | 0.390 | 0.496 | 0.640 | 1.439 | 0.282 |
| OR | 0.466 | 0.744 | 3.114 | 0.079 | 2.659 | 0.093 |
| CST | 2.995 | 0.134 | 5.074 | 0.022 | 2.691 | 0.093 |
| C_PHP | 0.405 | 0.762 | 2.022 | 0.198 | 1.524 | 0.273 |
| C_PH | 1.138 | 0.466 | 0.577 | 0.619 | 0.769 | 0.564 |
| C_FP | 2.792 | 0.138 | 4.361 | 0.032 | 2.956 | 0.077 |
| C_FPH | 0.109 | 0.897 | 0.607 | 0.619 | 0.191 | 0.902 |
| C_PO | 1.648 | 0.356 | 0.681 | 0.600 | 1.620 | 0.254 |
| F | 0.588 | 0.684 | 4.497 | 0.030 | 5.182 | 0.008 |
| CS_A | 1.510 | 0.372 | 0.058 | 0.943 | 2.250 | 0.129 |
| CS_P | 1.608 | 0.356 | 7.333 | 0.004 | 2.733 | 0.093 |
| CS_S | 0.684 | 0.646 | 0.712 | 0.600 | 1.497 | 0.273 |
| TR_A | 0.183 | 0.889 | 3.150 | 0.079 | 1.874 | 0.192 |
| TR_P | 1.071 | 0.477 | 5.560 | 0.016 | 2.169 | 0.136 |
| TR_S | 0.131 | 0.897 | 1.446 | 0.328 | 1.254 | 0.342 |
| CC_Body | 3.636 | 0.077 | 2.804 | 0.102 | 2.613 | 0.093 |
| CC_Major | 0.854 | 0.568 | 3.623 | 0.061 | 2.569 | 0.093 |
| CC_Minor | 1.705 | 0.356 | 3.357 | 0.070 | 3.111 | 0.067 |
| CC_Tap | 0.217 | 0.888 | 1.023 | 0.460 | 0.545 | 0.673 |

**Table S15**. Sensitivity analysis on the impact of including socioeconomic status (SES) as additional covariate on the relationships between baseline tract-BAG measures (with age-bias correction) and the cumulative number of KSADS-5 diagnoses at baseline, at 2-y-follow-up, and the transition of the transdiagnostic status between baseline and 2-y-follow-up.

| Tract-based measure | $BAG_{Baseline}$ ~ $KSADS_{baseline}$ | | $BAG_{Baseline}$ ~ $KSADS_{followup}$ | | $BAG_{Baseline}$ ~ $KSADS_{conversion}$ | |
|---|---|---|---|---|---|---|
| | F | $p_{FDR}$ | F | $p_{FDR}$ | F | $p_{FDR}$ |
| Association Development Mode | 15.216 | < 0.001 | 11.817 | < 0.001 | 10.641 | < 0.001 |
| Subcortical/Limbic Development Mode | 6.238 | 0.008 | 6.986 | 0.005 | 4.770 | 0.012 |
| Whole-brain | 7.083 | 0.004 | 6.902 | 0.005 | 6.086 | 0.002 |
| AF | 9.180 | 0.001 | 4.193 | 0.037 | 7.096 | 0.001 |
| FAT | 9.844 | 0.001 | 8.669 | 0.002 | 7.323 | 0.001 |
| PAT | 14.004 | < 0.001 | 8.410 | 0.002 | 8.657 | < 0.001 |
| SLF2 | 8.252 | 0.002 | 5.305 | 0.016 | 4.316 | 0.014 |
| SLF3 | 5.160 | 0.018 | 2.173 | 0.166 | 4.462 | 0.014 |
| SLF1 | 7.523 | 0.003 | 8.574 | 0.002 | 8.073 | < 0.001 |
| IFOF | 5.533 | 0.014 | 3.742 | 0.051 | 2.716 | 0.077 |
| ILF | 3.143 | 0.092 | 1.734 | 0.246 | 0.877 | 0.482 |
| UF | 1.279 | 0.387 | 0.651 | 0.556 | 0.886 | 0.482 |
| MdLF | 3.911 | 0.053 | 2.415 | 0.143 | 4.367 | 0.014 |
| VOF | 1.732 | 0.283 | 0.822 | 0.502 | 1.828 | 0.187 |
| OR | 0.600 | 0.650 | 2.858 | 0.097 | 2.400 | 0.100 |
| CST | 3.688 | 0.058 | 5.356 | 0.016 | 3.107 | 0.054 |
| C_PHP | 0.518 | 0.681 | 2.184 | 0.166 | 1.653 | 0.224 |
| C_PH | 0.932 | 0.504 | 0.958 | 0.461 | 0.955 | 0.472 |
| C_FP | 3.683 | 0.058 | 4.730 | 0.025 | 3.539 | 0.037 |
| C_FPH | 0.330 | 0.767 | 0.618 | 0.556 | 0.320 | 0.811 |
| C_PO | 2.231 | 0.181 | 1.014 | 0.461 | 1.826 | 0.187 |
| F | 1.170 | 0.414 | 4.676 | 0.025 | 4.633 | 0.012 |
| CS_A | 2.275 | 0.181 | 0.278 | 0.757 | 2.469 | 0.096 |
| CS_P | 2.541 | 0.153 | 7.842 | 0.003 | 3.204 | 0.051 |
| CS_S | 0.374 | 0.759 | 0.719 | 0.538 | 1.274 | 0.346 |
| TR_A | 0.733 | 0.591 | 3.361 | 0.065 | 2.050 | 0.152 |
| TR_P | 1.614 | 0.304 | 6.103 | 0.009 | 2.600 | 0.085 |
| TR_S | 0.212 | 0.835 | 0.943 | 0.461 | 1.064 | 0.430 |
| CC_Body | 4.036 | 0.052 | 3.216 | 0.071 | 2.963 | 0.061 |
| CC_Major | 1.297 | 0.387 | 4.026 | 0.041 | 2.922 | 0.061 |
| CC_Minor | 2.511 | 0.153 | 3.541 | 0.058 | 3.480 | 0.037 |
| CC_Tap | 0.003 | 0.997 | 1.615 | 0.265 | 0.389 | 0.786 |

**Table S16. Scanning parameters of T1w images in HCP-D, ABCD and HBN datasets.**

**(a)**

| | HCP-D | ABCD | | |
|---|---|---|---|---|
| Scanner | Siemens Prisma | Siemens Prisma | GE 750 | Philips Achieva |
| Sequence | Multi-echo T1w MPRAGE | MPRAGE | | |
| TR/TE (ms) | 2500/(1.8/3.6/5.4/7.2) | 2500/2.88 | 2500/2 | 6.31/2.9 |
| TI (ms) | 1000 | 1060 | 1060 | 1060 |
| Flip angle (°) | 8 | 8 | 8 | 8 |
| FOV | 256 × 240 × 166 | 256 × 256 | 256 × 256 | 256 × 240 |
| Resolution (mm) | 0.8 × 0.8 × 0.8 | 1.0 × 1.0 × 1.0 | 1.0 × 1.0 × 1.0 | 1.0 × 1.0 × 1.0 |

**(b)**

| Site | RUBIC | CBIC | CCNY |
|---|---|---|---|
| Scanner | Siemens Tim Trio | Siemens Prisma | Siemens Prisma |
| Sequence | | MPRAGE | |
| TR/TE (ms) | | 2500/3.15 | |
| TI (ms) | | 1060 | |
| Flip angle (°) | | 8 | |
| %FOV Phase | | 100% | |
| Resolution (mm) | | 0.8 × 0.8 × 0.8 | |

Abbreviation: RUBIC: Rutgers University Brain Imaging Center; CBIC: CitiGroup Cornell Brain Imaging Center; CUNY: City College of New York

**Table S17. Scanning parameters of dMRI images in HCP-D, ABCD and HBN datasets.**

**(a)**

| Dataset | HCP-D | ABCD | | |
|---|---|---|---|---|
| Scanner | Siemens 3T Prisma | Siemens 3T Prisma | GE 3T 750 | Philips 3T Achieva |
| Sequence | Multiband EPI | | HARDI | |
| TR/TE (ms) | 3230/89.20 | 4100/88 | 4100/81.9 | 5300/89 |
| Flip angle (°) | 78 | 90 | 78 | 77 |
| b-values ($s/mm^2$) | 1500 (186-dirs)<br>3000 (184-dirs) | | 500 (6-dirs)<br>1000 (15-dirs)<br>2000 (15-dirs)<br>3000 (60-dirs) | |
| FOV | 210 × 210 | 240 × 240 | 240 × 240 | 240 × 240 |
| Resolution (mm) | 1.5 × 1.5 × 1.5 | 1.7 × 1.7 × 1.7 | 1.7 × 1.7 × 1.7 | 1.7 × 1.7 × 1.7 |
| Multiband acceleration | 4 | 3 | 3 | 3 |

**(b)**

| Site | RUBIC | CBIC | CCNY |
|---|---|---|---|
| Scanner | Siemens Tim Trio | Siemens Prisma | Siemens Prisma |
| TR/TE (ms) | | 3320/100.2 | |
| Flip angle (°) | | 90 | |
| b-values ($s/mm^2$) | | 1000 (64-dirs)<br>2000 (64-dirs) | |
| %FOV phase | | 100% | |
| Resolution (mm) | | 1.8 × 1.8 × 1.8 | |
| Multiband acceleration | | 3 | |

Abbreviation: RUBIC: Rutgers University Brain Imaging Center; CBIC: CitiGroup Cornell Brain Imaging Center; CUNY: City College of New York

**Table S18**. Neurocognitive measures and the corresponding abbreviations in this study

| Test | Subscale (measure) | Abbreviation | ABCD field |
|---|---|---|---|
| NIH Toolbox cognition battery[2,3] | Picture Vocabulary Test Score | nihtbx_pic_vocabulary | nihtbx_picvocab_uncorrected |
| | Flanker Test Score | nihtbx_flanker | nihtbx_flanker_uncorrected |
| | Dimensional Change Card Sort Test Score | nihtbx_card_sort | nihtbx_cardsort_uncorrected |
| | List Sorting Working Memory | nihtbx_list_sort | nihtbx_list_uncorrected |
| | Oral Reading Recognition Test Score | nihtbx_oral_reading | nihtbx_reading_uncorrected |
| | Pattern Comparison Processing Speed | nihtbx_pattern_comp | nihtbx_pattern_uncorrected |
| | Picture Sequence Memory Test Score | nihtbx_pic_seq_memory | nihtbx_picvocab_uncorrected |
| | Fluid Intelligence | nihtbx_fluidcomp | nihtbx_fluidcomp_uncorrected |
| | Crystal Intelligence | nihtbx_cryst_comp | nihtbx_cryst_uncorrected |
| | Total Intelligence | nihtbx_total_comp | nihtbx_totalcomp_uncorrected |
| Rey Auditory Verbal Learning Test (RAVLT) | Accuracy in the immediate test | verbal_learning_immediate | pea_ravlt_sd_trial_vi_tc |
| | Accuracy in the delayed test | verbal_learning_delayed | pea_ravlt_ld_trial_vii_tc |
| Cash choice task | Choice of short Smaller-sooner | cash_choice_prefer_ss | cash_choice_prefer_ss |
| | Choice of short Larger-later | cash_choice_prefer_ll | cash_choice_prefer_ll |
| Wechsler Intelligence Scale for Children-V (WISC-V) | Matrix Reasoning Test Scaled Score | matrix_reasoning | pea_wiscv_tss |
| Little Man Task | Efficiency ratio | little_man_efficiency | lmt_scr_efficiency |
| Recognition memory test | d’ | recog_memory | |
| fMRI tasks | Emotional n-back (% accuracy on 2-back - % accuracy on 0-back) | 2_back | tfmri_nb_all_beh_c2b_rate |
| | Stop Signal Reaction Time, mean estimation | SST_ssrt | tfmri_sst_all_beh_total_mssrt |
| | Monetary Incentive Delay (Mean earnings) | MID_total_earning | tfmri_mid_all_beh_t_earnings |

**Table S19**. Psychopathology-related measures and the corresponding abbreviations in this study

| Questionnaire | Subscale (measure) | Abbreviation | ABCD field (ABCD file) |
|---|---|---|---|
| Prodromal Questionnaire Brief Version (PQ-B)[4] | Severity score of prodromal psychosis | PQB_psychosis_severity | pps_y_ss_severity_score |
| Ten-item Mania Scale[5] | Total score of mania | PGBI_mania_total | pgbi_p_ss_score |
| Children Behavior Check List (CBCL)[6] | Anxious/Depressive symptoms | CBCL_syn_anxdep | cbcl_scr_syn_anxdep_r |
| | Withdrawn/Depressed symptoms | CBCL_syn_withdep | cbcl_scr_syn_withdep_r |
| | Somatic Complaints | CBCL_syn_somatic | cbcl_scr_dsm5_somaticpr_r |
| | Social Problems | CBCL_syn_social_prob | cbcl_scr_syn_social_r |
| | Thought Problems | CBCL_syn_thought_prob | cbcl_scr_syn_thought_r |
| | Attention Problems | CBCL_syn_attention_prob | cbcl_scr_syn_attention_r |
| | Rule-Breaking Behavior | CBCL_syn_rulebreak | cbcl_scr_syn_rulebreak_r |
| | Aggressive Behavior | CBCL_syn_aggresive | cbcl_scr_syn_aggressive_r |
| | Internalizing Problems | CBCL_internal | cbcl_scr_syn_internal_r |
| | Externalizing Problems | CBCL_external | cbcl_scr_syn_external_r |
| | Total Problems | CBCL_total | cbcl_scr_syn_totprob_r |
| | Depressive Problems | CBCL_dsm_depressive | cbcl_scr_dsm5_depress_r |
| | Anxiety Problems | CBCL_dsm_anxiety | cbcl_scr_dsm5_anxdisord_r |
| | Somatic Problems | CBCL_dsm_somatic | cbcl_scr_dsm5_somaticpr_r |
| | Attention-deficit/hyperactivity Disorder | CBCL_dsm_ADHD | cbcl_scr_dsm5_adhd_r |
| | Oppositional Defiant Problems | CBCL_dsm_opposit | cbcl_scr_dsm5_opposit_r |
| | Conduct Problems | CBCL_dsm_conduct_prob | cbcl_scr_dsm5_conduct_r |
| | Sluggish Cognitive Tempo | CBCL_SCT | cbcl_scr_07_sct_r |
| | Obsessive-Compulsive Problems | CBCL_OCD | cbcl_scr_07_ocd_r |
| | Stress Problems | CBCL_stress | cbcl_scr_07_stress_r |
| Urgency, Perseverance, Premeditation and Sensation seeking (UPPS-P)[7,8] | Lack of Planning | UPPS_lack_planing | upps_y_ss_lack_of_planning |
| | Sensation Seeking | UPPS_sensation_seeking | upps_y_ss_sensation_seeking |
| | Positive Urgency | UPPS_positive_urgency | upps_y_ss_positive_urgency |
| | Negative Urgency | UPPS_negative_urgency | upps_y_ss_negative_urgency |
| | Lack of Perseverance | UPPS_lack_perserverance | upps_y_ss_lack_of_perseverance |
| Behavioral Inhibition & Behavioral Activation Scales (BIS/BAS)[9] | Drive | BISBAS_drive | mh_y_bisbas |
| | Reward responsiveness | BISBAS_reward_resp | bis_y_ss_bas_rr |
| | Fun seeking | BISBAS_fun_seek | bis_y_ss_bas_fs |
| | BIS total score | BISBAS_bis_total | bis_y_ss_bis_sum |

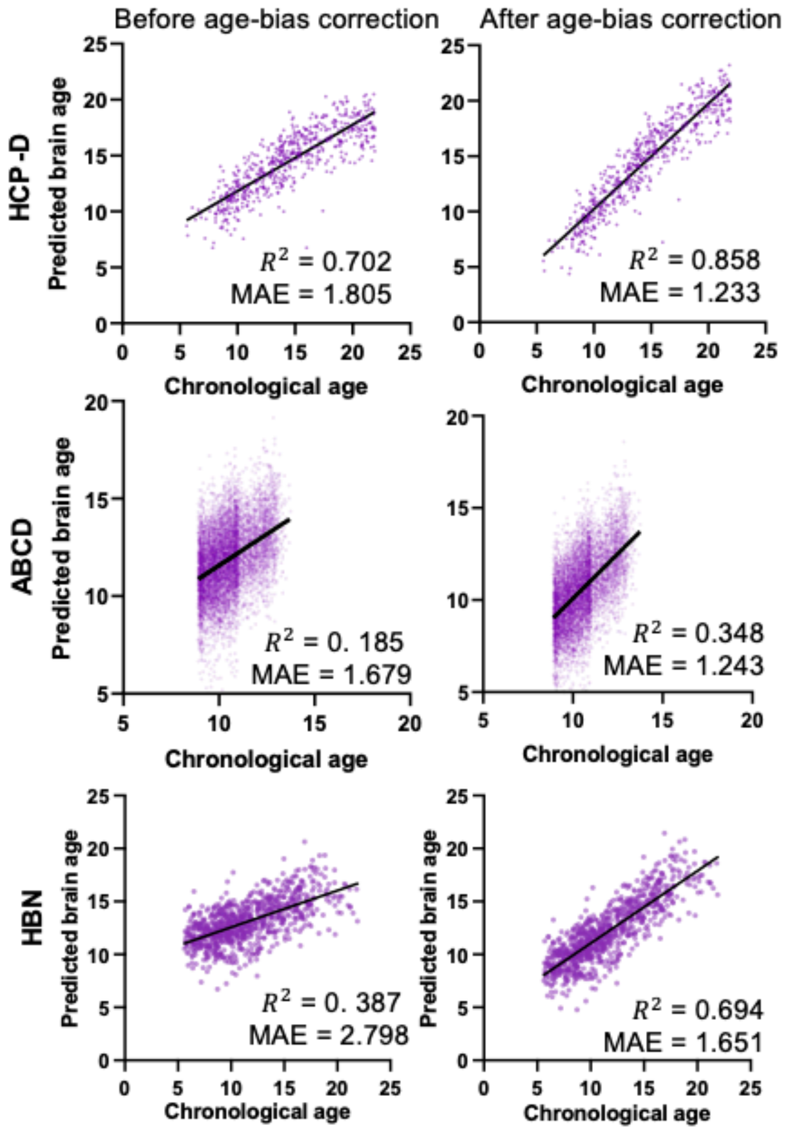


**Figure S1. Associations between the age predicted by whole-brain tracts and chronological age in cross-validation and independent testing.** Scatter plots depict the relationships between predicted brain age and chronological age using whole-brain tract-profiles from the Human Connectome Project in Development (HCP-D), Adolescent Brain Cognitive Development (ABCD) and Healthy Brain Network (HBN) pediatric mental health studies, respectively.

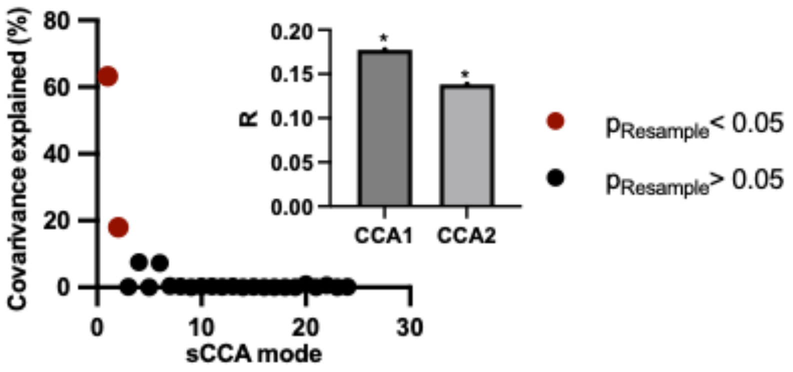


**Figure S2.** The first two canonical variates were statistically significant, as determined by permutation testing with false discovery rate (FDR) correction ($p < 0.05$).

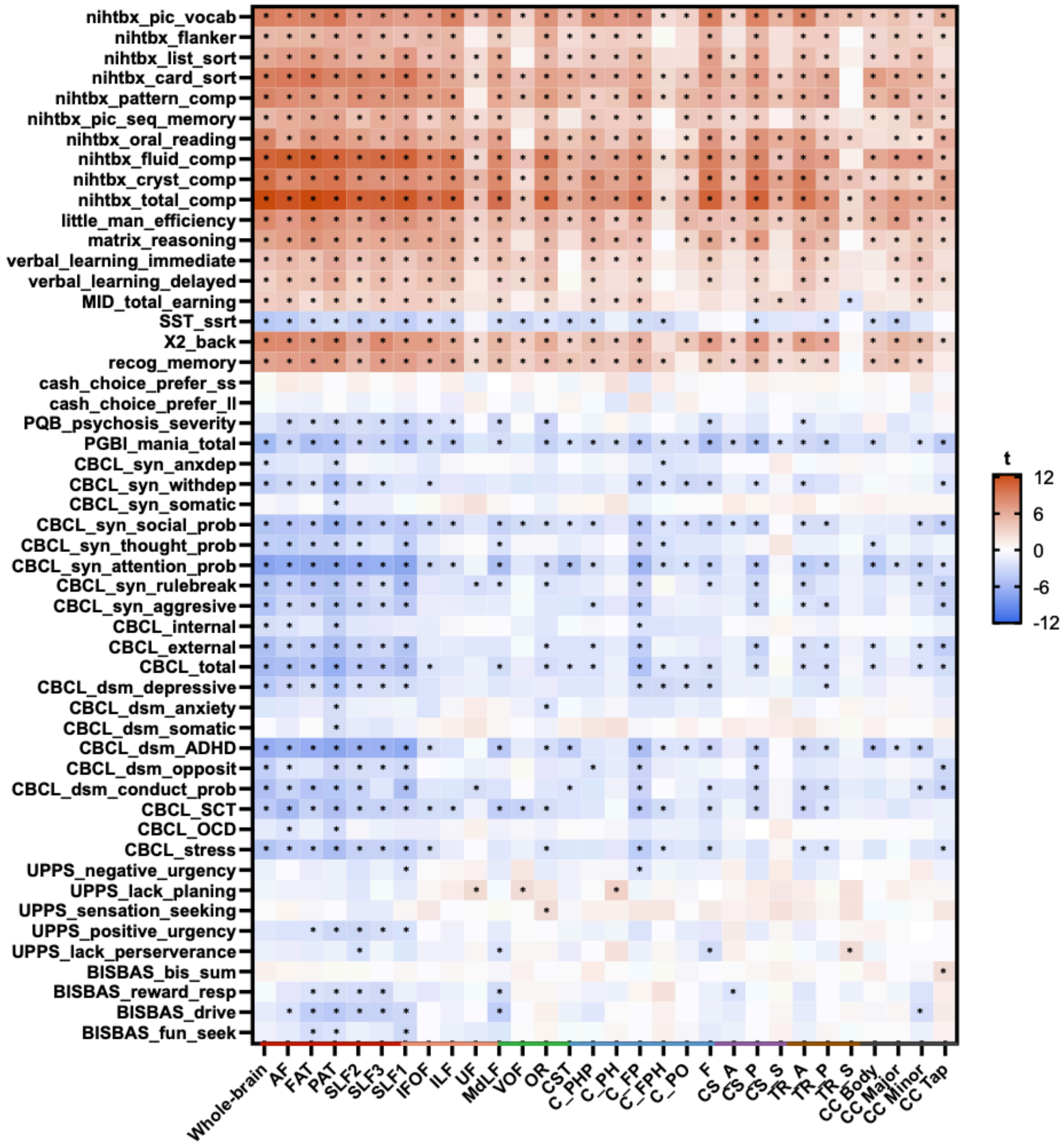


**Figure S3**. Heatmap of univariate associations between tract-based BAGs (with age-bias correction) and behavioral measures. The t-values from the Generalized Linear Model (GLM) analysis are shown. Asterisks indicate $p < 0.05$ after False Discovery Rate (FDR) correction. Abbreviation: BAG, Brain Age Gap.

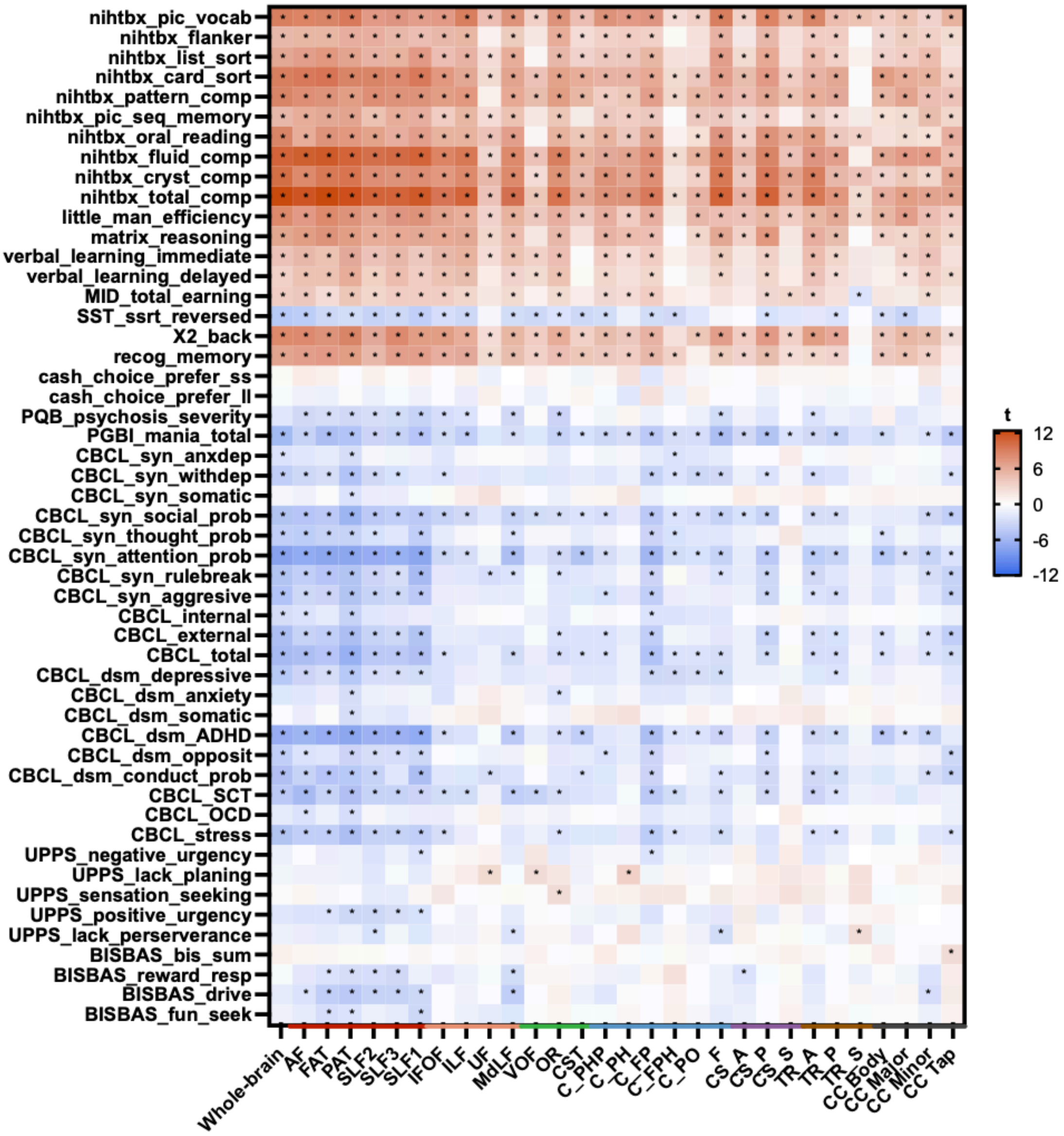


**Figure S4**. Heatmap of univariate associations between tract-based BAGs (without age-bias correction) and behavioral measures. The t-values from the Generalized Linear Model (GLM) analysis are shown. Asterisks indicate $p < 0.05$ after False Discovery Rate (FDR) correction. Abbreviation: BAG, Brain Age Gap.

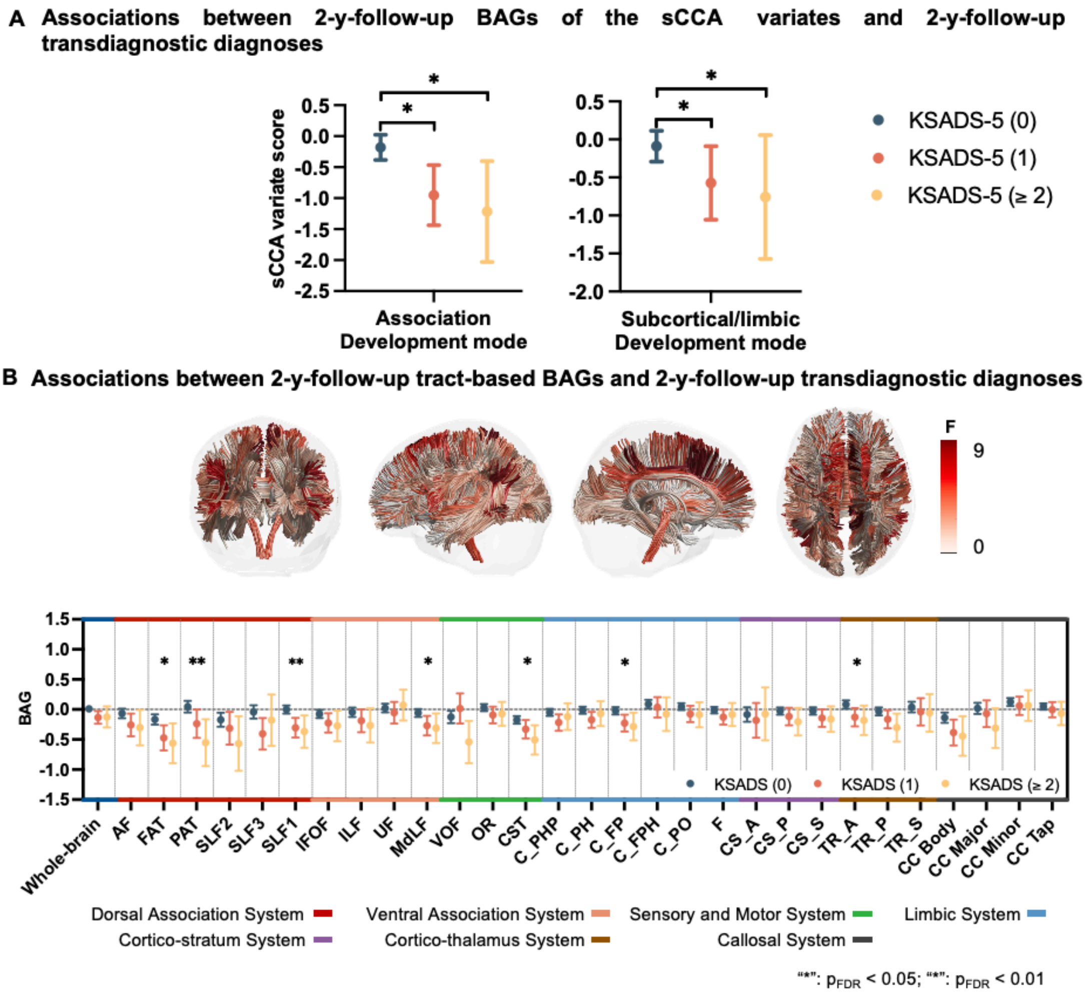


**Figure S5**. Relationships between 2-y-follow-up tract-BAG measures and 2-y-follow-up transdiagnostic status. Group-wise comparisons were conducted by GLM analyses. (A) Groupwise comparison of baseline sparse canonical correlation analysis (sCCA) variate scores with their baseline cumulative numbers of psychiatric disorders. (B) Associations between baseline tract-based BAGs and the cumulative number of 2-y-follow-up cumulative Kiddie Schedule for Affective Disorders and Schizophrenia for DSM-5 (KSADS-5) diagnoses. In the upper panel, the F-values of group-wise KSADS-5 effects for different tracts are plotted. The bottom panel shows the group-wise comparison results of tract-based BAGs in groups with 0, 1 or >=2 current diagnoses. Error bars represent 95% confidence intervals. "*" $p_{FDR} < 0.05$; . "**" $p_{FDR} < 0.01$. See Table S1 for abbreviations of the tracts. Abbreviations: BAG, brain-age gap; FDR, false discovery rate.

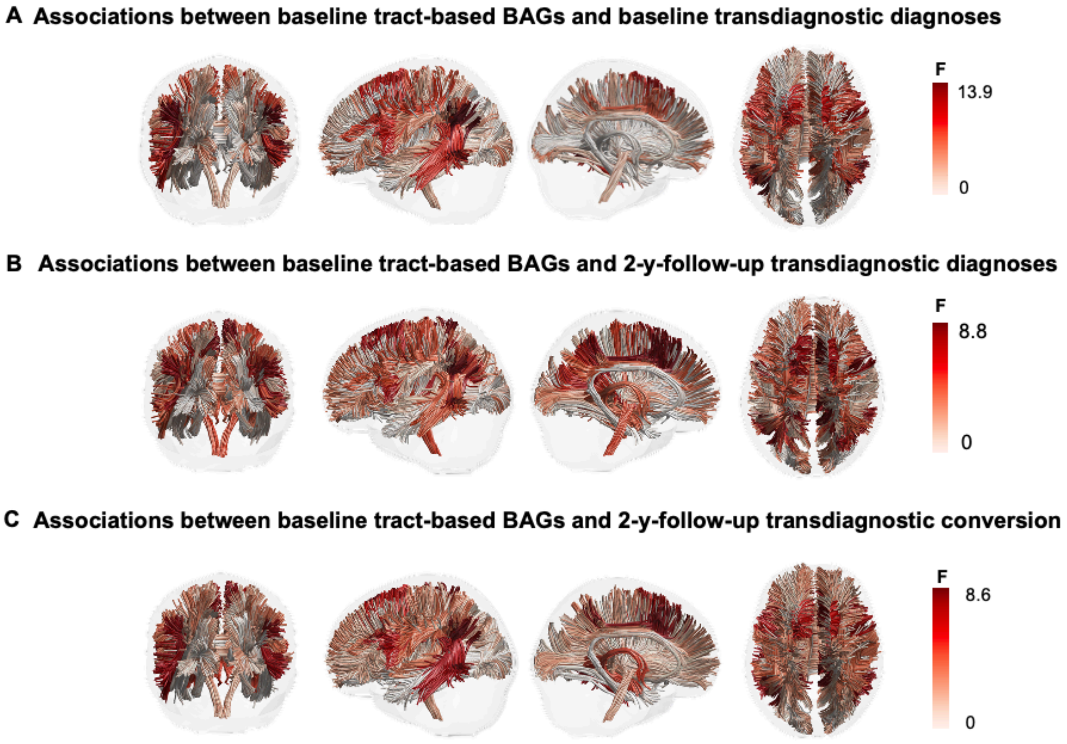


**Figure S6**. Relationships between baseline tract-BAGs and and the cumulative number of KSADS-5 diagnoses at baseline (A), at 2-y-follow-up (B), and the transition of the transdiagnostic status between baseline and 2-y-follow-up (C). In panels A and B, the F values of group-wise KSADS-5 effects for different tracts are plotted. The color of each tract in the upper panel corresponds to the F value of the tract-based BAGs. The panel C displays the specific tract-based BAGs of four groups with different transdiagnostic diagnosis status. Abbreviations: BAG, brain-age gap; KSADS-5: Kiddie Schedule for Affective Disorders and Schizophrenia for DSM-5.

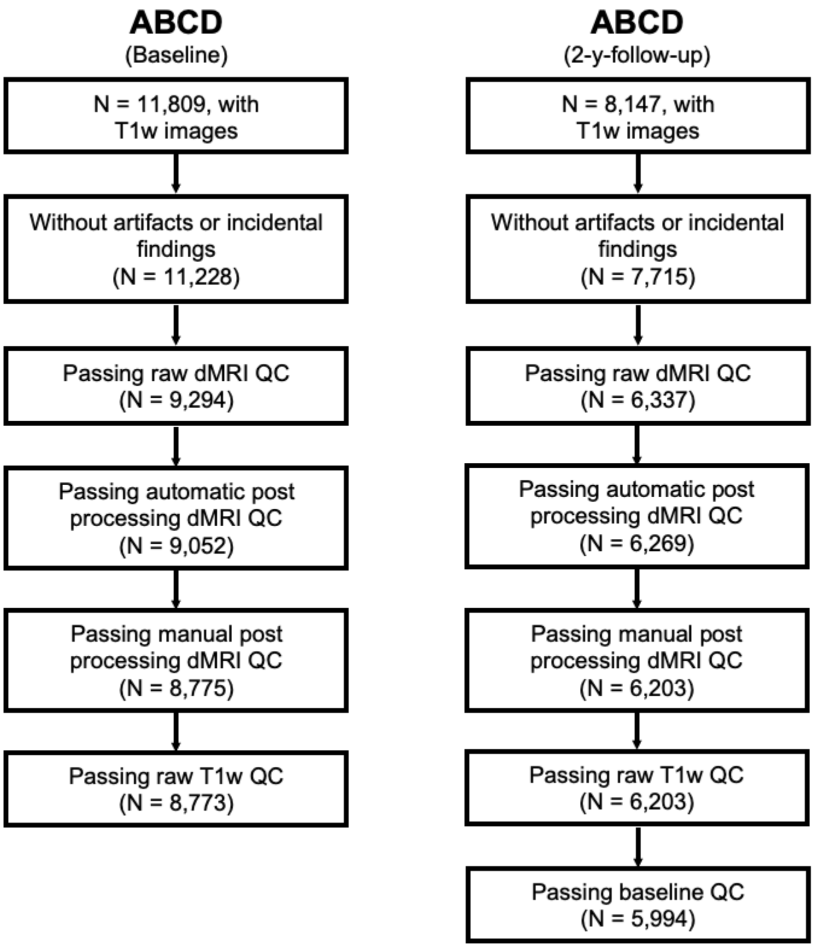


**Figure S7. Quality control (QC) procedure for the ABCD dataset at baseline and 2-y-follow-up.**

**Reference**


1. Desikan, R.S., *et al.* An automated labeling system for subdividing the human cerebral cortex on MRI scans into gyral based regions of interest. *Neuroimage* **31**, 968-980 (2006).
2. Luciana, M., *et al.* Adolescent neurocognitive development and impacts of substance use: Overview of the adolescent brain cognitive development (ABCD) baseline neurocognition battery. *Dev Cogn Neurosci* **32**, 67-79 (2018).
3. Rosenberg, M.D., *et al.* Behavioral and Neural Signatures of Working Memory in Childhood. *J Neurosci* **40**, 5090-5104 (2020).
4. Loewy, R.L., Pearson, R., Vinogradov, S., Bearden, C.E. & Cannon, T.D. Psychosis risk screening with the Prodromal Questionnaire — Brief Version (PQ-B). *Schizophrenia Research* **129**, 42-46 (2011).
5. Youngstrom, E.A., Frazier, T.W., Demeter, C., Calabrese, J.R. & Findling, R.L. Developing a 10-item mania scale from the Parent General Behavior Inventory for children and adolescents. *J Clin Psychiatry* **69**, 831-839 (2008).
6. Achenbach, T.M. & Verhulst, F. Achenbach system of empirically based assessment (ASEBA). *Burlington, Vermont* (2010).
7. Zapolski, T.C., Stairs, A.M., Settles, R.F., Combs, J.L. & Smith, G.T. The measurement of dispositions to rash action in children. *Assessment* **17**, 116-125 (2010).
8. Barch, D.M., *et al.* Demographic, physical and mental health assessments in the adolescent brain and cognitive development study: Rationale and description. *Dev Cogn Neurosci* **32**, 55-66 (2018).
9. Pagliaccio, D., *et al.* Revising the BIS/BAS Scale to study development: Measurement invariance and normative effects of age and sex from childhood through adulthood. *Psychol Assess* **28**, 429-442 (2016).
